# Dynamic changes in gas solubility of xylem sap reiterate the enigma of plant water transport under negative pressure

**DOI:** 10.1101/2022.01.06.475193

**Authors:** Luciano Pereira, Steven Jansen, Marcela T. Miranda, Vinícius S. Pacheco, Lucian Kaack, Gabriel S. Pires, Xinyi Guan, Juliana L.S. Mayer, Eduardo C. Machado, H. Jochen Schenk, Rafael V. Ribeiro

**Affiliations:** Laboratory of Plant Physiology “Coaracy M. Franco”, Center R&D in Ecophysiology and Biophysics, Agronomic Institute (IAC), Campinas SP, Brazil; Institute of Systematic Botany and Ecology, Ulm University, Albert-Einstein-Allee 11, 89081 Ulm, Germany; Laboratory of Crop Physiology, Department of Plant Biology, Institute of Biology, P.O. Box 6109, University of Campinas (UNICAMP), 13083-970, Campinas, SP, Brazil; Laboratory of Plant Anatomy, Department of Plant Biology, Institute of Biology, P.O. Box 6109, University of Campinas (UNICAMP), 13083-970, Campinas, SP, Brazil; Department of Biological Science, California State University Fullerton, 800 N. State College Blvd., CA 92831-3599 Fullerton, USA

## Abstract

Despite a long research history, we do not fully understand why plants are able to transport xylem sap under negative pressure without constant failure. Microbubble formation via direct gas entry is assumed to cause hydraulic failure, while the concentration of gas dissolved in xylem sap is traditionally supposed to be constant, following Henry’s law. Here, the concentration of soluble gas in xylem sap was estimated *in vivo* using well-watered *Citrus* plants under varying levels of air temperature and photoperiodic exposure, and compared to modelled data. The gas concentration in xylem sap showed non-equilibrium curves, with a minimum over- or undersaturation of 5% compared to gas solubility based on Henry’s law. A similar diurnal pattern was obtained from the gas concentration in the cut-open conduits and discharge tube, and oversolubility was strongly associated with decreasing xylem water potentials during transpiration. Although our model did not explain the daily changes in gas solubility for an anisobaric situation, oversolubility characterises nanoconfined liquids, such as sap inside cell walls. Thus, plants are able to transport sap under negative pressure with relatively high amounts of dissolved gas, providing them with a buffering capacity to prevent hydraulic failure, despite diurnal changes in pressure and temperature.

## Introduction

Although the flow of water through plants has utmost importance for the functioning of our biosphere, hydrology and agriculture, there are still open questions about the actual mechanisms behind water transport in plants (Jansen & Schenk, 2015). Long-distance transport in plants takes place in the xylem tissue, and represents a transpiration-driven process under negative pressure (Tyree & Zimmermann, 2002). This means that xylem sap is physically in a metastable state, and that water transport is prone to hydraulic failure due to embolism (Dixon & Joly, 1895). The water transport capacity of plants may indeed become reduced by large gas bubbles, with consequences for leaf gas exchange, plant water status, and eventually plant die-back (Choat *et al*., 2012; Anderegg *et al*., 2016). The water conducting xylem cells, which include multicellular vessels and unicellular tracheids, are known to become dysfunctional by air-entry under drought stress or freeze-thaw cycles. As there is increasing evidence that embolism formation in plants is relatively rare and limited to extreme drought events (Cochard & Delzon, 2013; Dietrich *et al*., 2019; Schuldt *et al*., 2020; Powers *et al*., 2020), one of the most important shortcomings in our understanding of water transport in plants concerns the long-standing question why xylem conduits are not constantly embolised at mild levels of drought stress or when the atmospheric demand is high. Transport of water under negative pressure is highly challenging and can be achieved in artificial systems, such as synthetic trees or a siphon with a height difference above 10 m, but only when pure and completely degassed water is used (Wheeler & Stroock, 2008; Boatwright *et al*., 2015). If water transport in xylem is not constantly interrupted by embolism, do plants have any unknown trick to prevent fast and frequent embolism formation?

There is clear evidence that embolism spreading during drought stress occurs through bordered pits, which represent openings in the secondary cell wall of conductive cells (Zimmermann, 1983; Sperry & Tyree, 1988; Kaack *et al*., 2019). Evidence that gas may cross cell walls and nucleate embolism was obtained by a gas pressurization technique (Cochard *et al*., 1992), in which a large pressure gradient is generated between the xylem sap and the gas surrounding the conduits. From a mechanistic point of view, the nanosized pores in the pit membranes are assumed to explain embolism spreading following the Young-Laplace equation (Tyree & Zimmermann, 2002). However, the complex and multicellular nature of xylem and the nanoscale processes at the gas-solid-water interface add complexity to the mechanisms underlying embolism formation and propagation. Moreover, the presence of both nanobubbles and amphiphilic lipids in xylem sap suggest that embolism formation does not rely on the dynamics of a single bubble crossing a pit membrane (Schenk *et al*., 2015, 2017; Kaack *et al*., 2019, 2021; Ingram *et al*., 2021).

Firstly, xylem sap is not pure water, and various surfactants present in sap may reduce its surface tension and therefore the internal xylem pressure for bubble formation (Schenk *et al*., 2017, 2021; Yang *et al*., 2020). Secondly, pit membranes are three-dimensional, thin, fibrous, and mesoporous media, with geometric interactions and deformation adding even more complexity to the gas-liquid interface as multiple pore constrictions exist along the gas pathway (Kaack *et al*., 2019, 2021; Zhang *et al*., 2020a). Thirdly, the concentration of gas dissolved in sap may be oversaturated during the warm periods (Schenk *et al*., 2016), with bark tissue and xylem cell walls acting as gas diffusion barriers (Wang *et al*., 2015). Oversaturation is a metastable state due to the energy requirement for bubble nucleation, which may, however, be overcome by mechanical stress or rapid changes in pressure and temperature (Mori *et al*., 1976). In fact, embolism formation by gas oversaturation was recently described as effervescence in gas pressurization experiments (Yin & Cai, 2018). Moreover, Henry’s law does not apply to the nanoscale dimensions of intervessel pit membranes and secondary walls because nanoconfined liquids contain higher amounts of dissolved gas, frequently referred to as oversolubility, as compared to the solubility predicted for a bulk liquid (Pera-Titus *et al*., 2009; Ho *et al*., 2015; Coasne & Farrusseng, 2019).

Because gas can diffuse through cell walls of conduits (Wang *et al*., 2015), the concentrations of atmospheric gases in both gaseous and liquid phases of xylem are assumed to be fairly close to the equilibrium at all times, i.e., that xylem sap is saturated following Henry’s law (Tyree & Zimmermann, 2002). However, little is known about gas solubility of xylem sap when embolised and sap-filled conduits are under anisobaric conditions (Mercury *et al*., 2003). It is known that gas solubility increases slightly with decreasing negative liquid pressure (Mercury *et al*., 2003; Lidon *et al*., 2018), and the effect of temperature should have a higher influence on gas solubility than pressure (Schenk *et al*., 2016). Based on membrane inlet spectrometry, Schenk et al. (2016) showed that xylem sap in *Distictis buccinatoria* was either saturated or supersaturated with atmospheric gases.

Here, we tested the unconventional hypothesis that the aqueous phase in xylem conduits (i.e., xylem sap) is indeed enriched with gases, and that gas concentrations may vary throughout the day due to changes in temperature, xylem water potential and in CO_2_ concentration due to respiration. We expected that gas solubility would increase during the day following a decrease in water potential. Gas solubility would also be affected by temperature and gas composition, as gases have different solubilities, and CO_2_/O_2_ concentrations are driven by respiration and photosynthesis (Hölttä *et al*., 2009). To test this hypothesis, we conducted a series of experiments in which we embolised cut-open vessels in xylem tissue of branches of well-irrigated plants. The technical challenge of measuring dissolved gas concentration in xylem of living plants was overcome by *in vivo* monitoring of gas discharge using a Pneumatron device (Pereira *et al*., 2016, 2020a), upgraded with a CO_2_ sensor. Data obtained was then compared with predicted data based on a gas diffusion model considering xylem anatomy and gas diffusion coefficients. The hypothesis that xylem sap can be oversaturated without embolism formation would indicate that xylem has a substantial buffering capacity to avoid constant hydraulic failure under moderate levels of drought, which is in line with solid and empirical evidence on the temporal frequency of embolism.

## Materials and Methods

### Plant material

*In vivo* experiments were performed with *ca*. 1-m tall orange trees [*Citrus sinensis* (L.) Osbeck grafted on *Citrus paradisi* Macf. x *Poncirus trifoliata* (L.) Raf.], with a basal stem diameter of *ca*. 15 mm. Plants were cultivated in pots (4.5 L) filled with substrate composed of pine bark and grown under greenhouse conditions, where air temperature varied from 18 to 42°C. Three to seven trees were used for each experiment, which were carried out inside a growth chamber (model PGR15, Conviron, Winnipeg MB, Canada). After acclimating plants to each condition, they were exposed to constant or variable temperature and variable duration of day light, depending on the experiment.

### Stem gas discharge measurements

A lateral branch was cut and connected to a Pneumatron device (Pereira *et al*., 2020a; Trabi *et al*., 2021) for gas discharge measurements (Fig. 1). The length of the cut size branch that was still attached to the main stem varied from 5 cm to 30 cm. Therefore, it was possible that vessels longer than 15 cm were directly connected to the main stem. In most experiments, the cut size branch had no leaves, but we also tested if the presence of leaves would affect the amount of gas extracted. We assumed that the cut-open vessels at the cut end of the branch were air-filled and under atmospheric pressure, as would be expected when cutting transpiring branches in air. However, the intact vessels that were connected to the cut-open vessels via intervessel pit membranes would still be filled with xylem sap as our plants were always well-irrigated. Vulnerability curves based on pneumatic measurements were conducted to show that embolism spreading from cut-open vessels to neighbouring intact vessels did not occur until a certain xylem water potential had been reached (Fig. S1), which was considerably lower than the water potential experienced by our irrigated plants. Therefore, native embolism in intact vessels and embolism spreading was not expected during our experiments. We did not aim to measure embolism resistance and our measurements were focused on gas diffusion at relatively high xylem water potentials.

**Fig. 1.**
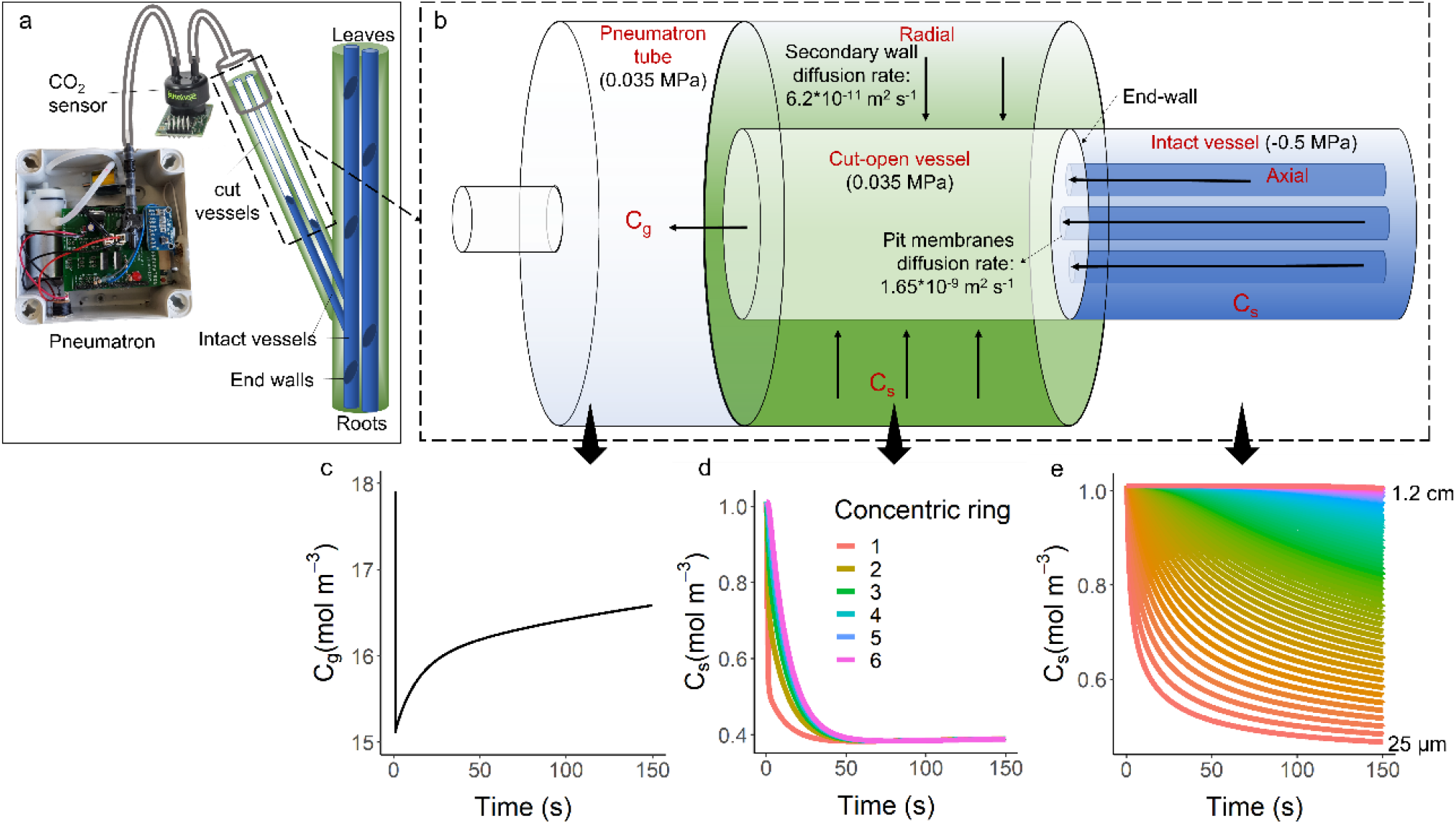
(a) Experimental set-up of an *in vivo* pneumatic experiment, showing the connection of the Pneumatron device to a cut branch of a living plant. While the cut-open conduits are air-filled and function as an extension of the discharge tube, intact vessels (i.e. vessels that are not cut open) remain filled with xylem sap under well-watered conditions; (b) scheme of the main parameters considered for the Sap Gas Extraction model, showing an intact and functional vessel (on the right) connected axially to a cut-open vessel via an end-wall and intervessel pit membranes. The cut-open and embolised vessel is connected radially to the surrounding tissues, from where the gas can be extracted and enter the Pneumatron tube (on the left). The diffusion through pit membranes is much higher than through lignified cell wall (10^−9^ *vs*. 2*10^−11^ m^2^ s^-1^, respectively). When a vacuum is applied to the cut-open vessel, the gas concentrations in xylem sap (C_s_) are modelled during 150 seconds of gas extraction at six concentric rings that surround the cut-open vessels (d) and at different distances from the pit membranes over time (e), which is also visualized by the small cylinders inside the intact vessel in b). The gas concentration (C_g_) in the embolised vessels and Pneumatron tubes is shown over time (c). Here, we considered a xylem water potential of −0.5 MPa and 25°C.

Radial gas diffusion through bark and lignified tissues is one or two orders of magnitude lower than axial gas diffusion through pit membranes (Sorz & Hietz, 2006; Yang *et al*., 2021). Gas may also be extracted from tissues surrounding the cut-open vessels, as suggested by Yang et al. (2021). The amount of Gas Discharged (GD) in moles was calculated considering the initial (P_i_) and final (P_f_) pressures, by using the ideal gas law:

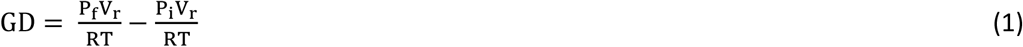

where R was the gas constant (8.3144621 J mol^-1^ K^-1^), V_r_ was the volume of the discharge tube, and T was the room temperature (in K).

Based on empirical and modelling evidence, we found previously that extraction of gas by diffusion from non-embolised vessels requires measuring intervals much longer than 15 seconds (Paligi *et al*., 2021; Yang *et al*., 2021). Therefore, we took gas discharge measurements over 150 seconds, with the initial pressure taken after 1 s, and the final pressure after 150 s. We were able to show that the amount of gas extracted did not come from embolised, intact conduits because this would have increased the total amount of gas discharged by one order of magnitude (Fig. S1).

The absolute amount of gas extracted (GD) varies with the sample size and/or anatomy. Thus, to facilitate comparison among *Citrus* trees, we considered the percentage of gas discharged (PGD), and calculated it from the maximum (GD_max_) and minimum (GD_min_) amount of gas extracted during measurements, as follows:

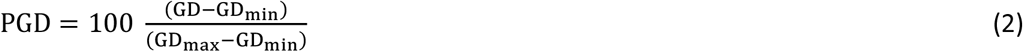

Similarly, values relative to the maximum GD (PGD_max_) were also used to highlight the total proportion of GD when necessary.

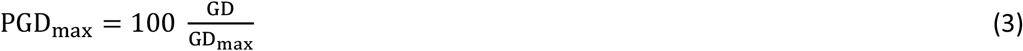

We conducted successive sets of experiments, firstly to understand the patterns of gas discharged, and then to identify traits correlated to it, as detailed below.

### Experiment 1: Gas discharge dynamics

In the first experiment, we evaluated the duration of gas extraction from the proximal side of a cut branch. We predicted that wound response would eventually occur and reduce gas diffusion on the cut end of the branch. For this, plants were kept under air temperature of 30°C and photosynthetic active radiation (PAR) of 400 μmol m^-2^ s^-1^ for 12 hours, and then subjected to darkness for another 12 hours. The measurements were stopped when the amount of gas extracted was constant and very low.

### Experiment 2: Varying light and temperature

Besides temperature – a key factor for gas solubility according to the Henry’s law – we also noticed in preliminary experiments that the amount of gas extracted from plants was strongly affected by light exposure. For this reason, the plants were exposed to varying temperature and light to evaluate their effects on the gas discharged from branches. While maintaining air temperature unchanged, cycles of 12 hours of light (PAR of 400 μmol m^-2^ s^-1^) and 12 hours of darkness were imposed to plants. After three days, plants were illuminated over 24 hours (entire day), while air temperature was changed from 15°C (12 hours) to 30°C (12 hours). The sequence of those treatments varied as follows: constant light/variable temperature followed by constant temperature/variable light. Using another set of plants, the opposite sequence was tested: constant temperature/variable light followed by constant light/variable temperature.

### Experiment 3: Xylem water potential, leaf gas exchange, and CO_2_ concentration inside branches

We sought for mechanistic or non-direct correlations that would help us to explain stem gas kinetics such as xylem water potential, leaf gas exchange, and CO_2_ concentration in the gas discharged from branches. Due to the technical challenge of measuring all these traits at the same time, especially leaf gas exchange, and considering the chronological development of our experiments, we did not measure all these traits simultaneously, but water potential was a common trait among all experiments.

Gas discharge measurements every 15 min were synchronised with measurements of xylem water potential using a stem psychrometer (PSY1, ICT International, Armidale NSW, Australia). The psychrometer was installed on the main stem according to the manufacturer instructions. CO_2_ assimilation (*A*_n_), stomatal conductance (*g*_S_), intercellular CO_2_ concentration (*C*_i_), and transpiration (*E*) were measured every 15 min on fully developed leaves with an infrared gas analyzer (Li-6400, Licor Inc., Lincoln NE, USA). During leaf gas exchange measurements, air CO_2_ concentration varied naturally, and light intensity and temperature followed the chamber configuration: air temperature was kept constant at 30°C to remove its possible effect on GD; the photoperiod was 12 hours under a PAR of 400 μmol m^-2^ s^-1^, followed by a dark period of 12 hours.

For quantifying the CO_2_ concentration in the gas discharged, a CO_2_ sensor (SprintIR-W-F-20, Gas Sensing Solutions, Cumbernauld, UK) was installed within the gas discharge tube and electronically connected to the Pneumatron so that the readings could be done in sync with the pressure sensor. The CO_2_ measurements were corrected by pressure considering a deviation of 1% kPa^-1^ according to the manufacturer instructions.

### Experiment 4: Removal of leaves and stopping stem photosynthesis

We investigated whether gas extraction was mechanistically correlated to water potential as described above. For this, we removed all leaves from the plants that were monitored after two or three days of gas discharge measurements, aiming to stop transpiration and water potential changes. We kept the plants under constant temperature (23°C) and a photoperiod of 16 hours (PAR of 400 μmol m^-2^ s^-1^). However, we noticed that water potential varied between light and dark conditions over various days, most likely due to gas exchange of green *Citrus* branches. To avoid this change in xylem water potential, we repeated the experiment by covering up the main stem and all branches with aluminium foil once all leaves had been removed.

### Experiment 5: Background gas extraction

We estimated the minimum and constant amount of gas extracted from the non-vessel surface of the cut branch, or from leakage in the tube connection. For this, we measured GD as described in experiment 1 for at least two days. Then, we cut the lateral branch keeping less than 2 cm connected to the Pneumatron, and sealed the proximal cut with a polyvinyl acetate glue. The gas extracted from this short stem segment was calculated as the percentage of maximum gas discharged (PGD_max_) before the branch was cut and considered to represent the background gas extracted from surrounding tissues, including intercellular spaces, parenchyma, and fibres. This percentage was used to confirm if the minimum amount of gas measured in experiment 1 represented only background gas or additional gas from the xylem. The lowest percentage, either from the experiment 1 or 5, was considered to be the background gas. This value was subtracted from the total amount of gas extracted from sap to estimate the gas concentration in sap following on a conservative and careful approach.

### Vessel length distribution and stem anatomy

We estimated vessel length distribution based on gas conductivity measurements using a Pneumatron, as gas conductivity is related to the amount of open vessels within a stem segment with a given length (Pereira *et al*., 2020b). Lateral branches of plants like those used in our experiments were evaluated. We also collected segments of the same branches for estimating the vessel diameter and vessel lumen fraction. For this, we cut transverse sections of fresh samples, which were imaged using a light microscope. At least 100 vessels were analyzed using the software ImageJ (version 1.52a; National Institutes of Health) and the equivalent circle diameter, the vessel lumen fraction (V_F_, %), and the fraction of a vessel pit field that is pit membrane (F_PM_, %) were calculated (Scholz *et al*., 2013).

Transmission electron microscopy (TEM) was applied to small wood samples from a *Citrus* plant growing at the botanical garden of Ulm University. Fresh material was prepared following a standard protocol, including fixation, dehydration, embedding, and sectioning, which was similar to Zhang *et al*. (2020b) and Kaack *et al*. (2021). The intervessel pit membrane thickness was measured based on TEM images, which were made with a JEOL 1400 TEM (JEOL, Tokyo, Japan) at an accelerating voltage of 120 kV, and with a digital camera (Soft Imaging System, Münster, Germany). The average pit membrane thickness was based on 43 intervessel pit membranes. Moreover, TEM observations were applied to determine the nature of the imperforate tracheary elements, and the potential occurrence of tracheids.

### Modelling gas extraction from sap

We developed a Gas Extraction model (SAX) for xylem sap (Box 1 and S1) based on the Unit Pipe Pneumatic model (Yang *et al*., 2021) to estimate how anatomical and environmental parameters would affect gas discharge from branches. As mentioned above, cut-open vessels were assumed to be connected to intact, water-filled vessels (Fig. 1a). The gas concentration in the sap of intact vessels was assumed to be saturated and in equilibrium with embolized vessels (axial connection) and with sap in surrounding tissue (radial connection). When the sap was under negative pressure during transpiration, gas diffusion increased axially through pit membranes and radially through secondary walls, and reached a higher gas concentration after equilibrium (Mercury *et al*., 2003; Schenk *et al*., 2016). When the Pneumatron created a subatmospheric pressure in the embolized vessel and discharge tube, gas diffusion occurred towards the cut-open vessels. The gas concentration in xylem sap decreased in the intact vessels, depending on the distance from the pit membrane (Fig. 1c), and in the tissue surrounding the cut-open vessels (Fig. 1d). Then, the gas concentration increased in the embolised, cut-open vessels and discharge tube (Fig. 1c). A list of all abbreviations, units and definitions can be found in Table 1, and all the parameters and equations are detailed in the Supporting Information (Methods S1).

**Box 1.**
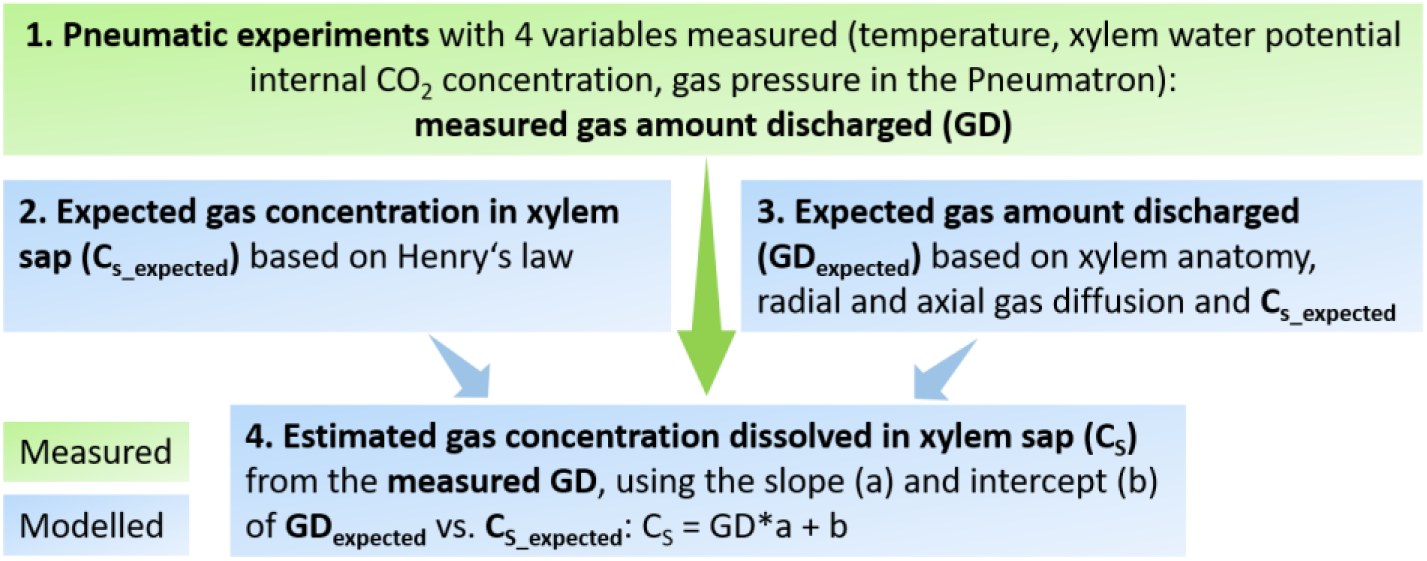
Simplified workflow of the gas extraction measurements and SAX-model, describing how the expected gas concentration in sap was compared to the estimated values from the gas discharge measured in plants of *Citrus sinensis*. A detailed workflow is shown in Box S1.

**Table 1.**
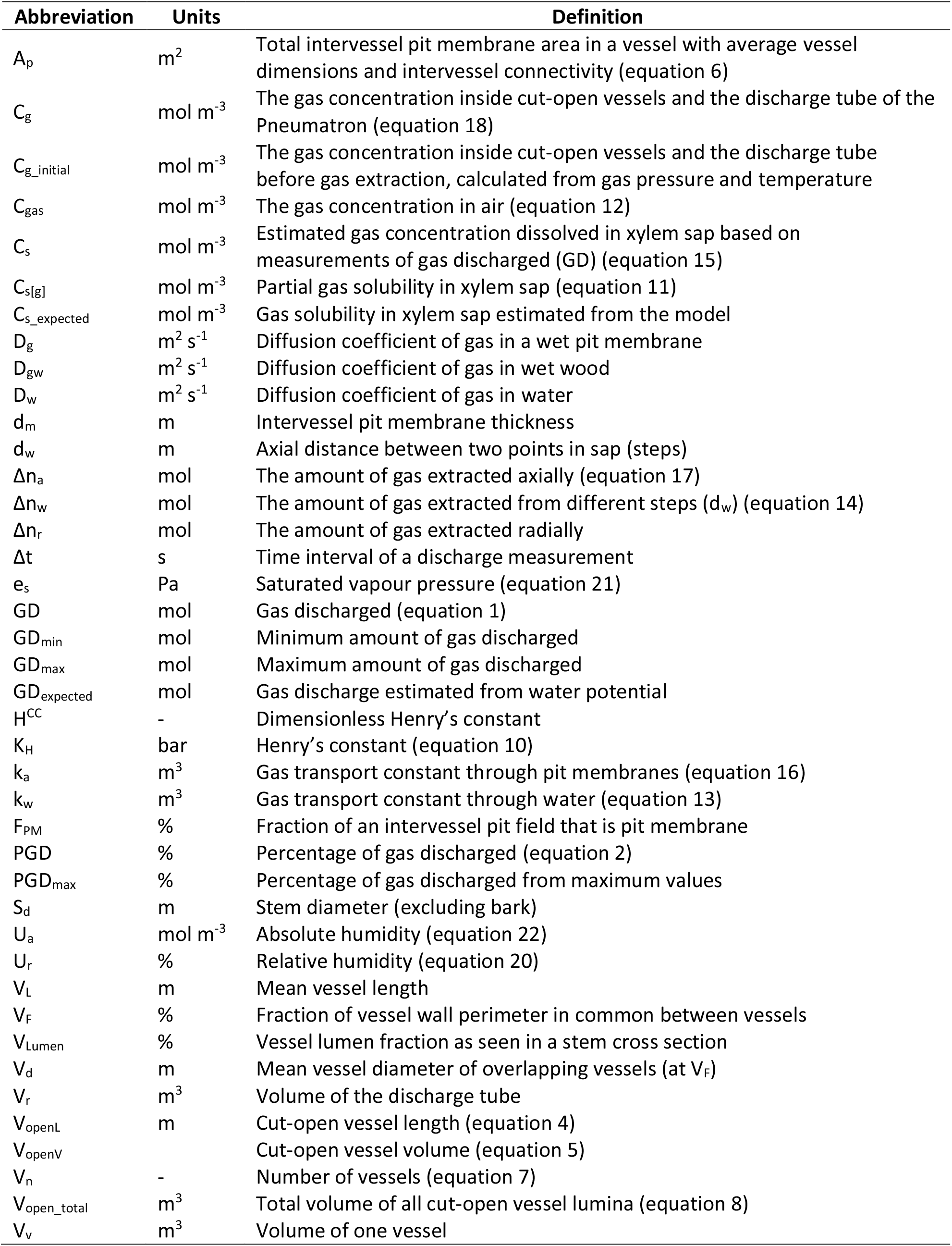
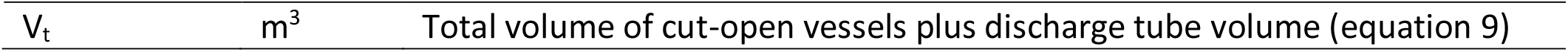
Overview of all abbreviations, with reference to their definitions and units, used for *in vivo* pneumatic measurements and a gas solubility model of xylem sap (equations in Methods S1).

| Abbreviation | Units | Definition |
| --- | --- | --- |
| $A_p$ | $m^2$ | Total intervessel pit membrane area in a vessel with average vessel dimensions and intervessel connectivity (equation 6) |
| $C_g$ | $mol\ m^{-3}$ | The gas concentration inside cut-open vessels and the discharge tube of the Pneumatron (equation 18) |
| $C_{g\_initial}$ | $mol\ m^{-3}$ | The gas concentration inside cut-open vessels and the discharge tube before gas extraction, calculated from gas pressure and temperature |
| $C_{gas}$ | $mol\ m^{-3}$ | The gas concentration in air (equation 12) |
| $C_s$ | $mol\ m^{-3}$ | Estimated gas concentration dissolved in xylem sap based on measurements of gas discharged (GD) (equation 15) |
| $C_{s[g]}$ | $mol\ m^{-3}$ | Partial gas solubility in xylem sap (equation 11) |
| $C_{s\_expected}$ | $mol\ m^{-3}$ | Gas solubility in xylem sap estimated from the model |
| $D_g$ | $m^2\ s^{-1}$ | Diffusion coefficient of gas in a wet pit membrane |
| $D_{gw}$ | $m^2\ s^{-1}$ | Diffusion coefficient of gas in wet wood |
| $D_w$ | $m^2\ s^{-1}$ | Diffusion coefficient of gas in water |
| $d_m$ | $m$ | Intervessel pit membrane thickness |
| $d_w$ | $m$ | Axial distance between two points in sap (steps) |
| $\Delta n_a$ | $mol$ | The amount of gas extracted axially (equation 17) |
| $\Delta n_w$ | $mol$ | The amount of gas extracted from different steps ($d_w$) (equation 14) |
| $\Delta n_r$ | $mol$ | The amount of gas extracted radially |
| $\Delta t$ | $s$ | Time interval of a discharge measurement |
| $e_s$ | $Pa$ | Saturated vapour pressure (equation 21) |
| GD | $mol$ | Gas discharged (equation 1) |
| $GD_{min}$ | $mol$ | Minimum amount of gas discharged |
| $GD_{max}$ | $mol$ | Maximum amount of gas discharged |
| $GD_{expected}$ | $mol$ | Gas discharge estimated from water potential |
| $H^{CC}$ | - | Dimensionless Henry's constant |
| $K_H$ | $bar$ | Henry's constant (equation 10) |
| $k_a$ | $m^3$ | Gas transport constant through pit membranes (equation 16) |
| $k_w$ | $m^3$ | Gas transport constant through water (equation 13) |
| $F_{PM}$ | % | Fraction of an intervessel pit field that is pit membrane |
| PGD | % | Percentage of gas discharged (equation 2) |
| $PGD_{max}$ | % | Percentage of gas discharged from maximum values |
| $S_d$ | $m$ | Stem diameter (excluding bark) |
| $U_a$ | $mol\ m^{-3}$ | Absolute humidity (equation 22) |
| $U_r$ | % | Relative humidity (equation 20) |
| $V_L$ | $m$ | Mean vessel length |
| $V_F$ | % | Fraction of vessel wall perimeter in common between vessels |
| $V_{Lumen}$ | % | Vessel lumen fraction as seen in a stem cross section |
| $V_d$ | $m$ | Mean vessel diameter of overlapping vessels (at $V_F$) |
| $V_r$ | $m^3$ | Volume of the discharge tube |
| $V_{openL}$ | $m$ | Cut-open vessel length (equation 4) |
| $V_{openV}$ | | Cut-open vessel volume (equation 5) |
| $V_n$ | - | Number of vessels (equation 7) |
| $V_{open\_total}$ | $m^3$ | Total volume of all cut-open vessel lumina (equation 8) |
| $V_v$ | $m^3$ | Volume of one vessel |
| $V_t$ | $m^3$ | Total volume of cut-open vessels plus discharge tube volume (equation 9) |

## Results

### Gas extraction over time under variable light and/or temperature

The difference between the amount of gas discharged under dark and light conditions (experiment 1) was on average 3.26*10^−6^ ± 1.39*10^−6^ mol, with a higher amount of gas extraction under light than under darkness. This difference between light and darkness became small after 4-5 days, reaching a minimum of 11.2 ± 1% after 13 days (Figs. 2 and S3).

**Fig. 2.**
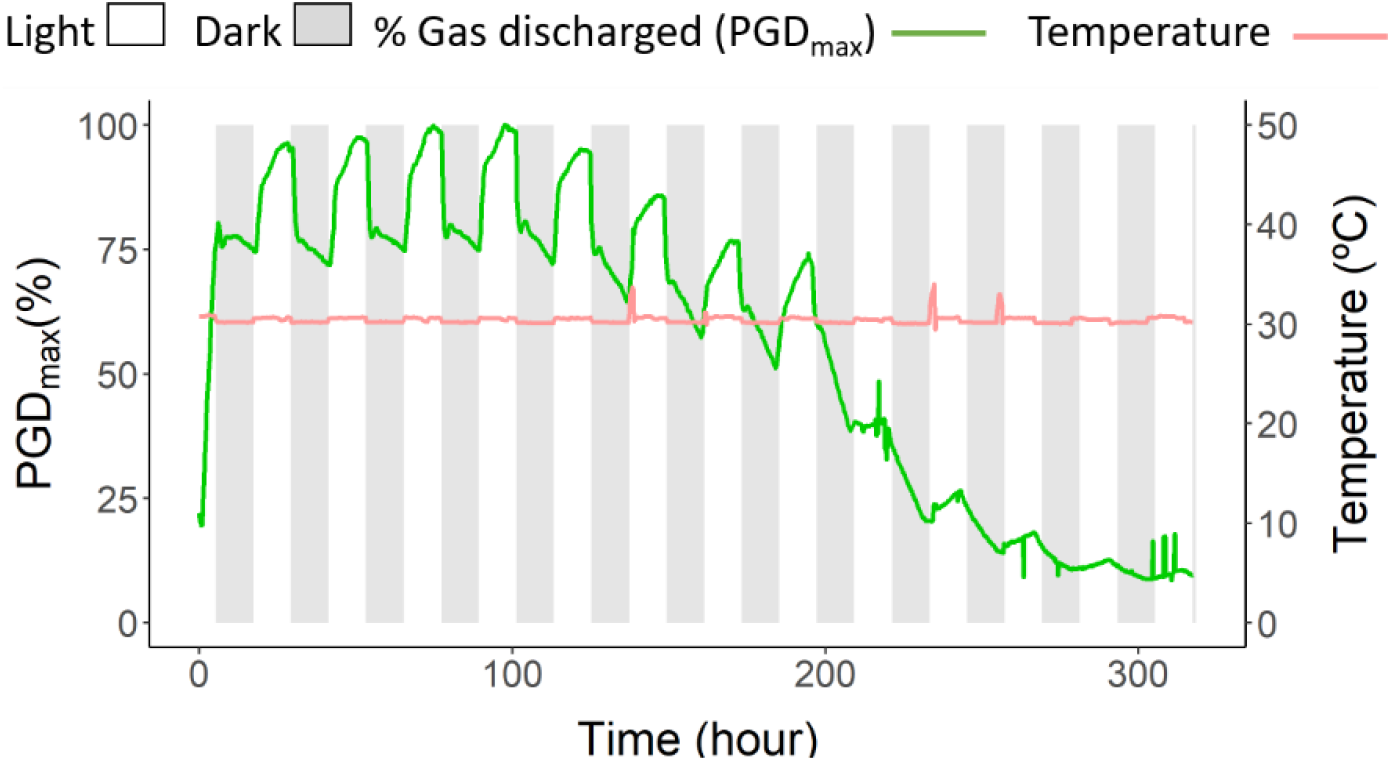
Temporal dynamics of the percentage of gas discharged related to the maximum value measured (PGD_max_, green line) for a cut side branch of a well-irrigated orange tree under 12 hours of light (PAR 400 μmol m^-2^ s^-1^, white bars) and 12 hours of darkness (grey bars) over 13 days inside a growth chamber (experiment 1). Air temperature is shown by the orange line. In Figure S3, all *Citrus sinensis* plants measured are shown.

Both temperature and light availability (experiment 2) were significantly correlated with the percentage of gas discharged (PGD), but the photoperiodic effect on PGD was larger than the temperature effect (Figs. 3 and S4). The sequence of the treatments (continuous light/variable temperature followed by continuous temperature/variable light *vs*. continuous temperature/variable light followed by continuous light/variable temperature) had no impact on the gas amount discharged (Fig. 3).

**Fig. 3.**
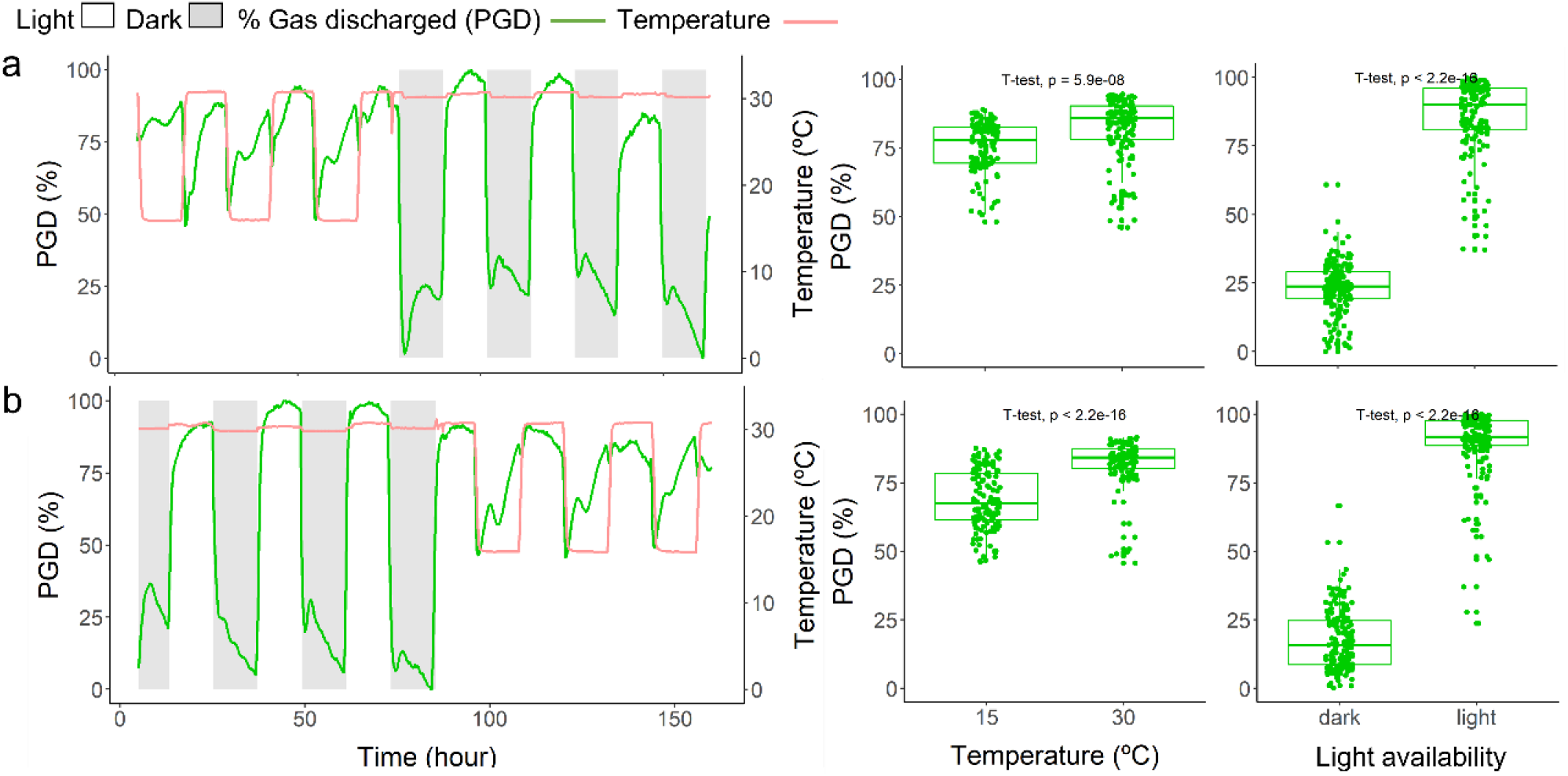
Temporal dynamics (a, b) of the percentage of gas discharged (PGD, green line) from a cut side branch of a well-irrigated orange tree under varying light or air temperature (experiment 2). Air temperature (orange line) varied between 15°C and 30°C, and light conditions between PAR 0 (grey bars) and 400 (white bars) μmol m^-2^ s^-1^. Box plots compare all data points from the plant shown in (a) or (b). A single plant was used for the experiment shown in a, and another plant was used for the experiment shown in b. Replications are shown in Figure S4.

### Is gas extraction related to xylem water potential, leaf gas exchange, intercellular CO_2_ concentration, or gas from background tissue?

PGD was strongly related to the xylem water potential (R=-0.88±0.05, Figs. 4 and S5, experiment 3) and the slope of a linear regression between these two variables was 123±30 % MPa^-1^, although the intercept presented larger variation (−7.2±33.9 %, Fig. 4). The relationship between xylem water potential and PGD was largely linear, except for a steep decline in PGD at water potentials close to 0 MPa.

**Fig. 4.**
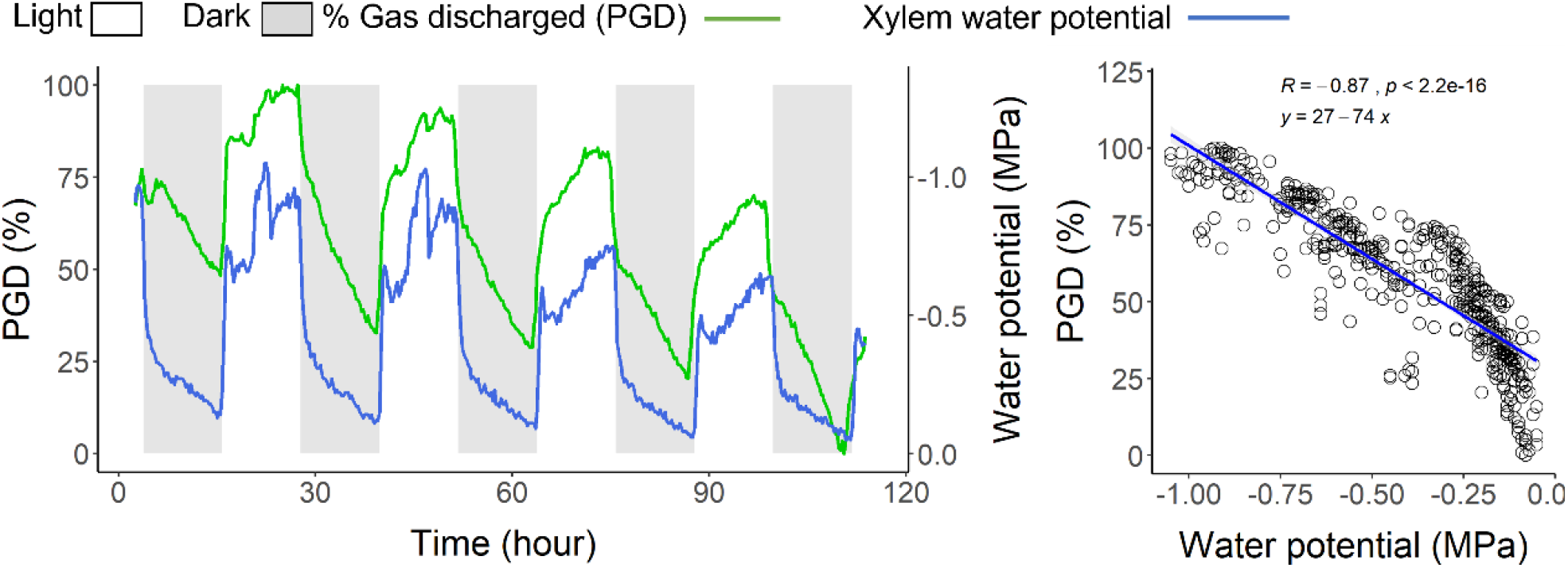
(a) Temporal dynamics of the percentage of gas discharged (PGD, green line) and xylem water potential (blue line) in a representative orange tree under 12 hours of light (white bars) and darkness (grey bars) over 4.5 days (experiment 3). In (b), PGD is correlated to the xylem water potential, using data shown in (a). All plants are shown separately in Figure S5.

The strong correlation between water potential and gas discharged remained when all leaves were removed, and the branches were covered up with aluminium foil (experiment 4). Under these conditions, water potential and PDG hardly changed between the 16 hours of light and 8 hours of darkness (Figs. 5 and S6). Interestingly, the CO_2_ concentration in the gas extracted responded to light conditions, with the highest CO_2_ concentrations found under darkness and being considerably lower during light periods (Fig. 5b).

**Fig. 5.**
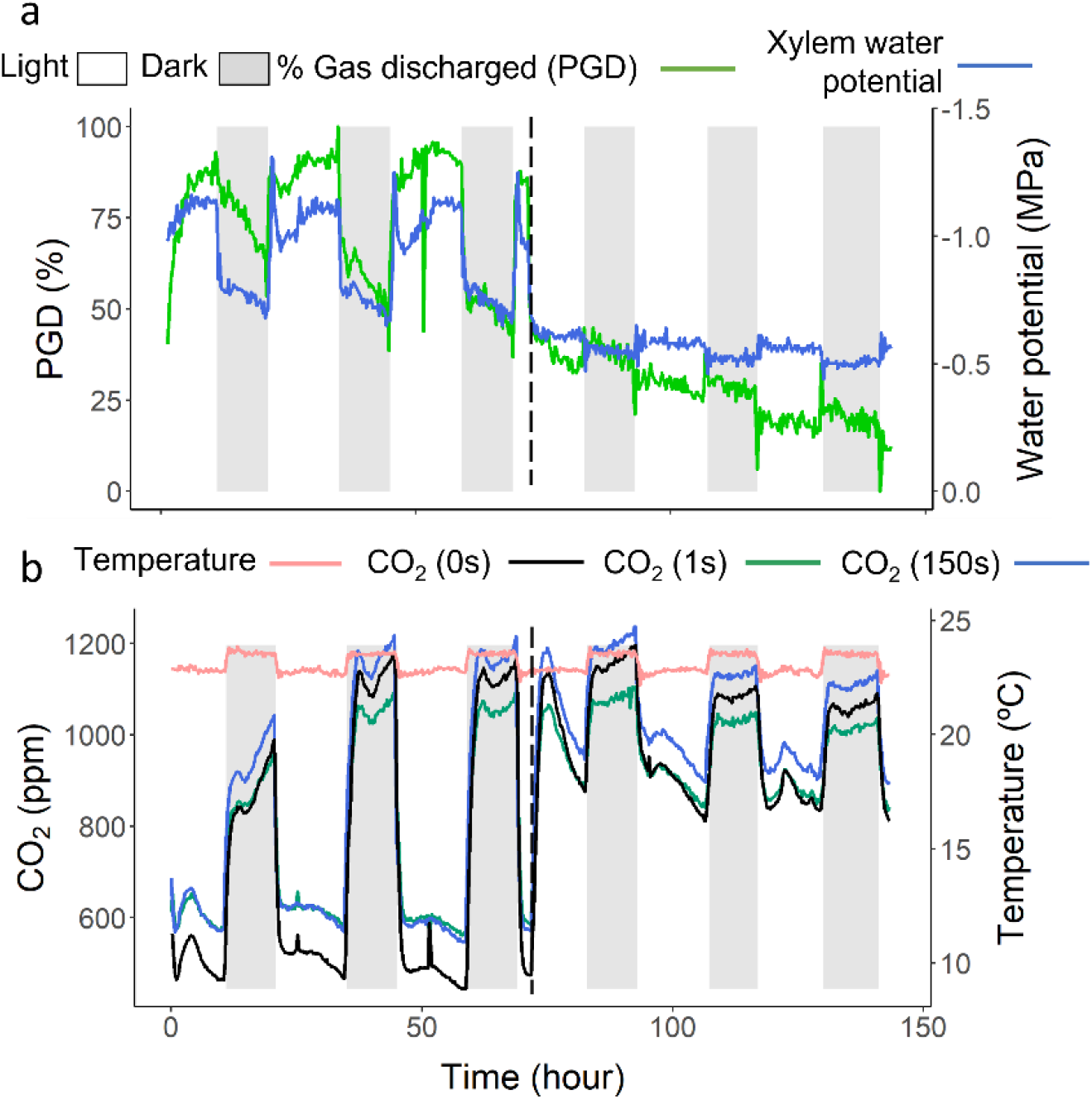
Temporal dynamics of (a) the percentage of gas discharged (PGD, green line), xylem water potential (blue line) and (b) CO_2_ concentration (green, blue and black lines) in a representative *Citrus* tree under 16 hours of light (white bars) followed by 8 hours of darkness (grey bars) over 6 days (experiments 3 and 4). CO_2_ concentration was measured before gas extraction started (black line), after the first extraction second (green line), and after 150 seconds of gas extraction (blue line). Air temperature dynamics is also shown in (b). All leaves were removed, and stems were covered with aluminium foil after 3 days (vertical dashed lines) to avoid changes due to transpiration and gas exchange as much as possible.

Leaf gas exchange was also significantly correlated with the amount of gas discharged (experiment 3), with a strong relationship for PGD *vs*. transpiration (R=0.78±0.12, Fig. S7) and PGD *vs*. CO_2_ assimilation (R=0.77±0.10, Fig. S8), and slightly weaker relationships for PGD *vs*. stomatal conductance and intercellular CO_2_ concentration (R=0.68±0.10 and −0.67±0.07, respectively, Figs. S9 and S10).

The gas extraction from the short stem segments (experiment 5) represented 28.7±9.5 % of the total gas discharge extracted from the lateral branches (Fig. S11), and was higher than the 11.2 ± 1% of gas extracted from lateral branches after 13 days (experiment 1).

### Modelling gas extraction from sap

The input values for the SAX model were either obtained from literature, or directly measured on the plants of *Citrus sinensis* (Table 2). Except for pit membrane thickness (d_m_), anatomical measurements were conducted on the same plants that were used in our experiments. For the modelling, we considered the mean values for the anatomical parameters (V_L_, V_F_, V_LUMEN_, V_d_, F_PM_, and d_m_, all measured and shown in Table 2 and Fig. S12).

**Table 2.**
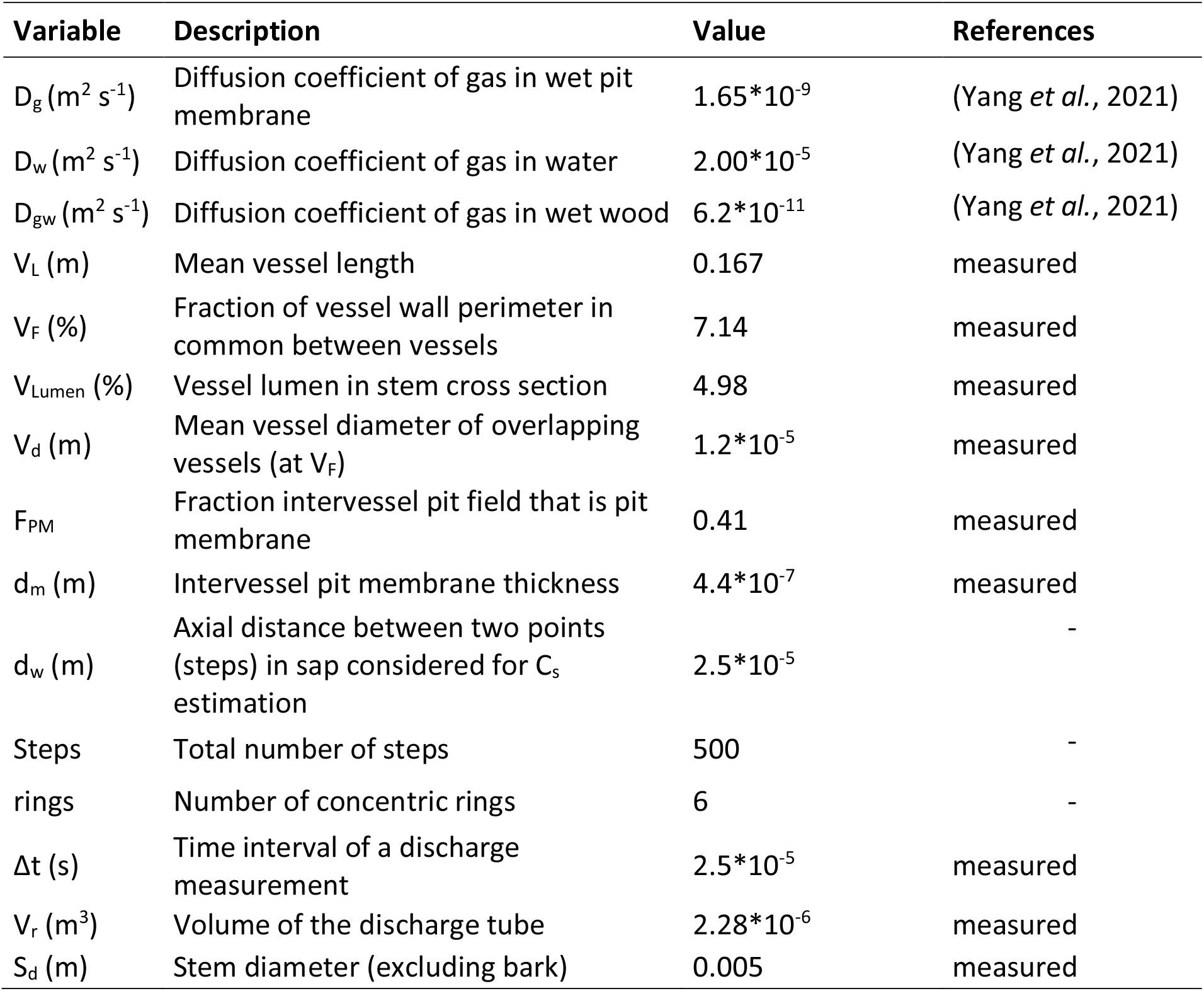
Variables used for modelling dissolved gas concentrations in xylem sap of *Citrus sinensis*.

The SAX model estimated the gas concentration in sap (C_s_) at least 5% above or below the expected gas concentration (C_s_expected_, Fig. 6a). On the other hand, when only the radial diffusion was considered, the model overestimated C_s_expected_ compared to C_s_, and explained less than 10% of C_s_. For the two model versions, we subtracted 11.2 % of gas from the total amount of gas extracted due to background gas (Figs. 2 and S11). Also, C_s_expected_ presented an opposite trend compared to the measured values of gas concentration in the discharge tube and cut-open vessel lumina (C_g_; Fig. 6b), and the measured gas discharge (PGD, Fig. 6b,c).

**Fig. 6.**
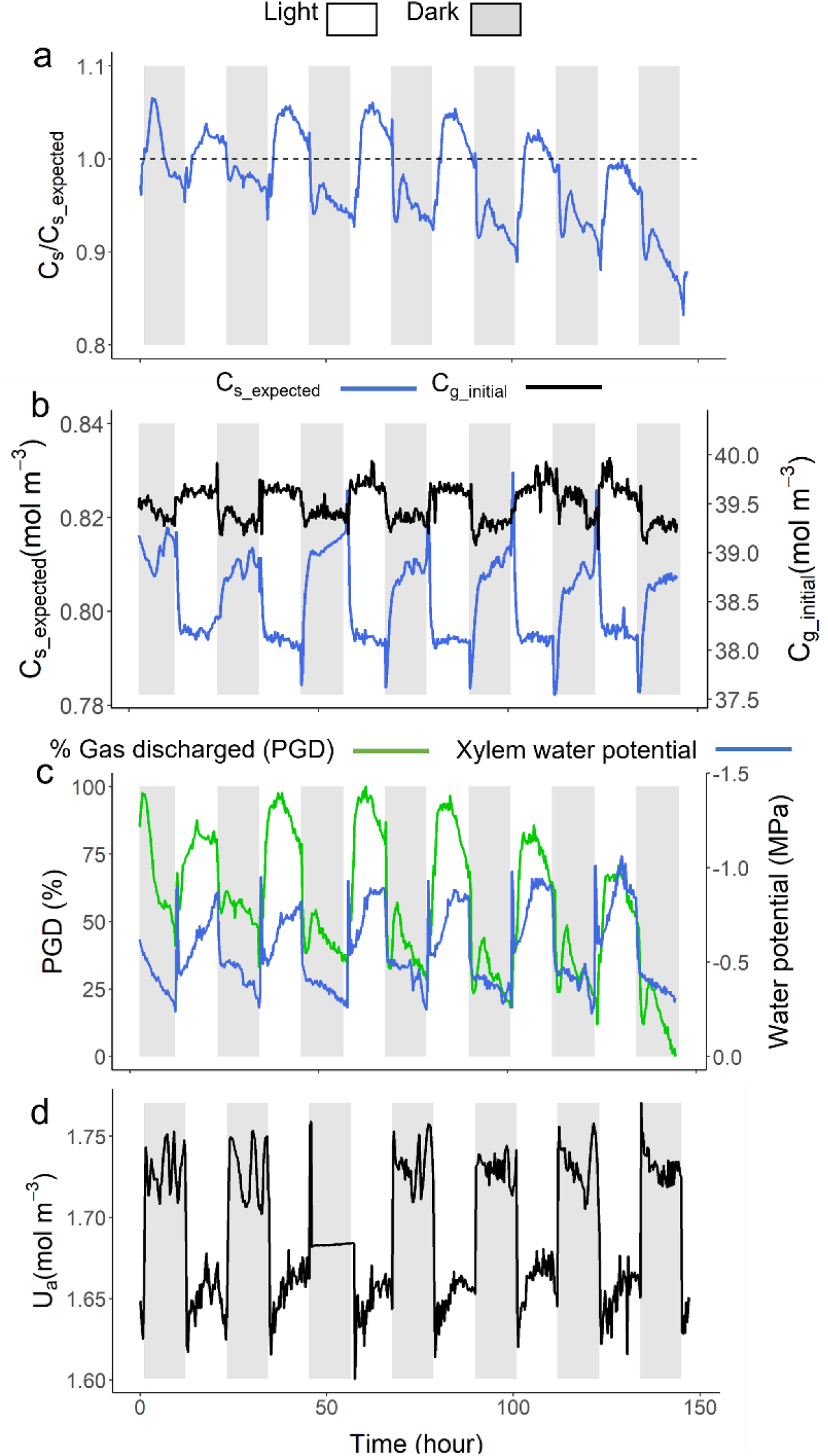
Temporal dynamics of the measured and modeled gas kinetics in a well-irrigated *Citrus sinensis* tree exposed to 12 hours of light (white bars) and 12 hours of darkness (grey bars). In (a), we show the gas concentration in xylem sap (C_s_) - estimated from the measured gas discharged - in relation to the expected gas concentration (C_s_expected_) - obtained from a model, when considering temperature, water potential and CO_2_ concentration. The dashed line represents the 1:1 line. In (b), C_s_expected_ (blue line) presents an opposite trend compared to the measured gas concentration (black line), and also to the gas discharge and water potential measurements, as shown in (c). It is also possible to see the opposite trend between C_s_ and the modelled absolute humidity of discharged air (in d).

The model presented some uncertainty for estimation of C_s_ related to the mean vessel diameter (V_d_), mean vessel length (V_L_), intervessel wall surface (V_f_), vessel lumen fraction (V_Lumen_), and the intervessel pit field fraction that is pit membrane (F_PM_). We tested for all possible combinations of maximum and minimum values for these traits, considering the standard deviations that were estimated from the samples measured (Fig. S13). As vessel length scales with vessel diameter (Cai *et al*., 2010; Liu *et al*., 2018; Yang *et al*., 2021), we assumed that there was a correspondence between the minimum and maximum values measured for both parameters, and that the intermediate values increased linearly. Moreover, the CO_2_ concentration (Fig. 5b) and the estimated absolute humidity (Fig. 6d) presented opposite trends to the gas discharge measured (Fig. 6c) and the gas concentration measured in the discharge tube and cut-open vessels (Fig. 6b), with the lowest and highest values of either CO_2_ concentration or absolute humidity being found under light and dark conditions, respectively (Figs. 5b and S6e). The CO_2_ concentration also increased during the gas extraction period (150 s), whether measured after 1 s or 150 s (green and blue lines), but usually presented a slightly higher concentration before the gas extraction started (Fig. S6, black lines). If the gas discharged were only composed of CO_2_ and water vapour, we would expect lower gas extraction volumes under light conditions, but higher gas extraction volumes during darkness.

## Discussion

Our results provide solid evidence that gas concentrations in xylem sap of *Citrus sinensis* plants were not constant, with values of at least 5% of the expected solubility (Fig. 6a) when plants were transpiring and under more negative water potentials (Fig. 4a). In this condition, the estimated versus expected gas concentrations in xylem sap deviated considerably from the 1:1 line (C_s_/C_s_expected_, Fig. 6a), showing a clear shortcoming of the estimated gas concentrations predicted by Henry’s law, even if the water potential, temperature, and CO_2_ concentration were considered. Under dark conditions, when there was little or no transpiration and the water potential of xylem sap was high, the gas concentration was probably undersaturated and below the 1:1 line. Because of the measurement deviation for some anatomical parameters, our SAX model included an uncertainty to estimate the exact gas concentration (Fig. S13). Interestingly, the model predicted oversolubility in six out of eight scenarios simulated. Most importantly, the measurements were variable and did not follow the Henry’s law, which raises the question on how plants are able to deal with dissolved gas concentrations in sap that deviate at least 5% from the expected equilibrium (Fig. 6a), and if changes in soluble gas concentration may affect bubble nucleation, and thus embolism.

### Xylem sap contains variable amounts of dissolved gas

Although temperature is negatively and exponentially related to gas solubility, and is theoretically its main determinant (Schenk *et al*., 2016), our *in vivo* measurements showed that temperature had a lower impact on gas concentration in xylem sap than changes induced by light (Fig. 3). The high gas concentrations estimated here by the SAX model agreed with the 1.1 ± 0.2 times higher concentration of dissolved gas in xylem sap of *Distictis buccinatoria* than what would be expected based on Henry’s law (Schenk *et al*., 2016). Additional evidence for high levels of gas dissolved in xylem sap is provided by the high gas concentration in the discharge tube and cut-open vessels prior to the pneumatic measurements (C_g_initial_), which also presented higher concentrations under light conditions. This was contrary to the expected concentration in xylem sap (Fig. 6b) and C_g_initial_ was measured independently before any gas was extracted, while considering the temperature and the gas pressure.

An alternative explanation for the increased gas extraction under light is that additional gas came from gas dissolved in the transpiration stream during an extraction time of 150 s because removing the leaves and excluding stem photosynthesis stopped the daily pattern in gas exchange. Interestingly, an increased amount of gas in xylem tissue during the day was also shown by Pate and Canny (1999), who extracted gas from root segments of *Xanthorrhea preissii*. Indirect evidence was also presented by Canny *et al*. (2007), who found the same daily pattern as described here and interpreted their results as daily cycles of embolism formation. However, this assumption remains controversial and is largely unsupported by recent literature (Wheeler *et al*., 2013; Martin-StPaul *et al*., 2017; Creek *et al*., 2020; Sorek *et al*., 2021). Moreover, since Pate and Canny (1999) extracted gas from non-transpiring root segments, the increased gas solubility during the day was probably not related to sap flow, reinforcing our hypothesis that plant xylem can deal with high gas concentrations dissolved in sap. Despite the conservative and critical approach followed here, which included accounting for gas extraction from background tissue, our experiments demonstrate that xylem sap can have a relative oversaturation. This finding supports the hypothesis that plant xylem can take up high amounts of dissolved gas (Pate & Canny, 1999; Schenk *et al*., 2016) without embolism formation.

After 4 to 5 days of gas discharge measurements, reduced gas concentrations were measured (Fig. 2), which may be due to a wound response. We speculate that clogging of xylem sap lipids or other substances occurred at the pit membranes (Schenk *et al*., 2021), which would increase the gas diffusion resistance, avoiding rapid changes in the equilibrium of gas concentration. A similar deposition of xylem sap lipids on or in pit membranes has been suggested over a growing season (Schmid & Machado, 1968; Sorek *et al*., 2021). As plants may face cut-open vessels due to herbivory or storm damage, the wound response to increase the gas diffusion resistance may also be a safe strategy to avoid embolism spreading (Guan *et al*., 2021), and could also prevent the entry of pathogens into xylem.

### Are there mechanistic explanations for the high gas concentrations in xylem sap?

The high gas concentrations of xylem sap cannot be explained by temperature, water potential, and CO_2_ concentration, and then additional mechanisms should be considered. Besides the fact that gas solubility increases slightly with decreasing negative liquid pressure (Mercury *et al*., 2003; Lidon *et al*., 2018), three interconnected and complementary hypotheses could explain gas oversolubility. First, xylem sap is known to be more complex than water, with amphiphilic, insoluble, and polar lipids providing a dynamic surface tension that is dependent on the local concentration of lipids per surface (Schenk *et al*., 2017; Yang *et al*., 2020). Lipid layers create diffusion barriers on gas-water interfaces (Borden & Longo, 2004), thereby potentially slowing equilibrium between the gas and liquid phase in xylem. Second, the mechanism behind the daily gas concentration pattern may be determined by the high cell wall resistance to gas diffusion, and the relatively low resistance through pit membranes (Fig. 1; (Sorz & Hietz, 2006; Yang *et al*., 2021)). This sap-confinement would result in a delay to reduce or enrich the gas concentration of xylem sap, changing from less to more saturated conditions, depending on the time of the day and the xylem water potential.

A third explanation is that oversaturation of gas is a well-known phenomenon of liquids in nanoconfined environments (Lidon *et al*. 2018; Coasne & Farrusseng 2019), which seems to be the case for xylem sap inside pit membranes and cell walls. Gas oversaturation in liquids, which is frequently described as oversolubility, can be explained either by gas adsorption or confinement, depending on the strength of the interaction between the gas-solid or liquid-solid interface, respectively (Ho *et al*. 2015; Coasne & Farrusseng 2019). The oversaturation effect is pronounced when pores are smaller than 15 nm, which is roughly the average pore size suggested for intervessel pit membranes (Choat *et al*., 2003, 2004; Kaack *et al*., 2019, 2021; Zhang *et al*., 2020b). Oversaturation of gas is even more relevant in pores that are a few nanometers in diameter (Miachon *et al*., 2008; Pera-Titus *et al*., 2009), which are present in the secondary cell walls of conduits (Donaldson *et al*., 2015, 2018). Oversaturation may also occur when a liquid is under negative pressure, such as at the liquid-gas interface in a nanopore that is exposed to dry air and consequently forms a concave meniscus (Lidon *et al*., 2018). This situation is similar to pores in a pit membrane that separate a functional from an embolized conduit (Fig. 1). Hence, physical interactions at the gas-solid-liquid-surfactant interface creates a complex, nanoscale system where Henry’s law cannot be applied (Pera-Titus *et al*., 2009). In this situation, liquids can be oversaturated up to 30 times compared to the value predicted for a bulk liquid, depending on the gas and solid species (Luzar & Bratko, 2005). It is unclear to what extent oversolubility of gas in nanoscale pores of cell walls and pit membranes affects the overall solubility of sap in conduit lumina, where confinement is at the microscale.

Our pneumatic experiments and SAX-model also showed that dissolved gas concentrations in xylem sap can be detected with the Pneumatron. In this case, the oversaturation measured may represent a mean bulk concentration of the gas dissolved in sap, including sap that is not inside pit membranes. Independently of the gas source, our data showed that dissolved gas is either over- or undersaturated in the aqueous phase in xylem and it is not equilibrated as expected by Henry’s law for an anisobaric situation. Although sources of non-dissolved gases may be present in fibres and intercellular spaces, where gas diffusion is 10^−4^ times faster than in water, the gas needs to be dissolved and to cross hydrated, secondary walls and liquids that surround the vessels. In terms of concentration, the predicted equilibrium between the dissolved and non-dissolved gas would decrease the gas concentration in both phases when gas is extracted by the Pneumatron. Thus, the non-dissolved gas source would contribute to the amount of gas extracted in a similar way as the dissolved gas. Extraction of non-dissolved gas could increase the distance from where gas can be extracted from the cut-open vessels and, hypothetically, increase the total amount of gas extracted radially. Therefore, our model could slightly overestimate gas solubility, although we expect a small impact because of (1) a higher radial than axial resistance, and (2) a relative small volume of fibre lumina and intercellular spaces in secondary xylem (Back, 1968; Gartner *et al*., 2004).

### How can plants transport sap that contain high concentrations of dissolved gas without constant embolism formation?

The density and surface tension of a liquid are dependent on many factors that could affect embolism spread through cell wall pores, traditionally described by the Young-Laplace equation (Schenk *et al*., 2015). Nanoconfined liquids in contact with solid surfaces present marked density oscillations (Coasne & Farrusseng, 2019), and the density as well as the surface tension may change with pressure, gas concentration, local compression or expansion of surfactants, and the curvature of the meniscus at the liquid-gas interface in a pore (Lidon *et al*., 2018). Hence, these parameters are dynamically adjusted and need to be taken in account to predict mass flow of gas across multiple pore constrictions of a pit membrane, which cannot be approached accurately by the Young-Laplace equation. Although some recent progress has been achieved in our understanding of the complex interactions between xylem sap, surfactants, pit membranes, and gases (Schenk *et al*., 2015, 2017, 2021; Pereira *et al*., 2018; Kaack *et al*., 2019, 2021; Yang *et al*., 2020; Zhang *et al*., 2020a; Guan *et al*., 2021), we currently are not able to precisely quantify the behaviour of these four elements under varying pressure.

Nevertheless, bubble nucleation may increase with gas concentration in liquids (Mori *et al*., 1976). Considering that xylem sap in pit membranes may be oversaturated with dissolved gas, rapid changes in pressure and temperature may nucleate bubbles since excessive volume and gaseous molecules dissolved in a liquid (V_g_) could then be released into vapour phase, as P_g_V_g_ = C_s_H^cc^V_g_ (Li *et al*., 2020), where P_g_ is the partial pressure of a given gas. This will also depend on the solid surface hydrophobicity, which can decrease the free energy and increase bubble nucleation. On the other hand, lipid-coated nanobubbles could be stable due to low and dynamic surface tension of lipid monolayers, which compensates the internal Laplace pressure (Schenk *et al*., 2015, 2017; Alheshibri *et al*., 2016; Ingram *et al*., 2021). Lipid-coated nanobubbles could allow higher gas concentration in xylem sap, probably buffering rapid changes. As lipid surfactants are present in sap and cover conduit surfaces and pit membranes (Schenk *et al*., 2017, 2018, 2021; Kaack *et al*., 2019), their functions should be explored in detail in further research on gas dynamics of xylem sap and embolism formation.

There is strong evidence that embolism spreads from an embolized vessel into a neighbouring one (Sano *et al*., 2011; Choat *et al*., 2016; Hochberg *et al*., 2016), which is usually explained by the air-seeding hypothesis (Tyree & Zimmermann, 2002). However, the presence of gas in contact with a water meniscus in pit membranes should increase gas diffusion, which would allow fast changes in gas concentration in the nano-confined sap and then induce bubble nucleation by excess gas concentration when the equilibrium between the liquid and gas phases is quickly unbalanced. However, fast changes in gas concentration are unlikely when water is present in a neighbouring vessel as the gas transport rate is decreased 10^−4^ times in liquid water. This would buffer changes in gas concentration under rapid pressure and temperature changes. Herein, we propose that mass flow of gas via air-seeding may not be the only mechanism behind embolism spreading since dynamic changes of gas concentration could play a role. The concentration of dissolved gas in xylem sap would especially be important when pressure and temperature are rapidly changing in the absence of water buffering the gas pathway through pit membranes. These rapid changes may explain why embolism is formed when xylem is cut under tension (Wheeler *et al*., 2013), and when xylem sap under negative pressure is shock-frozen (Pate & Canny, 1999; Umebayashi *et al*., 2016; Ogasa *et al*., 2019). As soluble gas concentration in sap increases under negative pressure (Pate & Canny, 1999; Schenk *et al*., 2016), a rapid change to a less negative pressure when the branch is cut would induce nucleation by the excessive volume of gas molecules dissolved in sap. However, if the negative pressure is progressively reduced, soluble gas may diffuse out of the sap and reach a new concentration equilibrium, without inducing embolism (Wheeler *et al*., 2013; Torres-Ruiz *et al*., 2015).

## Conclusion

The gas extraction method reported herein provides a straightforward and direct way to study temporal dynamics of gas concentration in plant xylem at a high resolution. Daily changes in gas solubility show undersaturation and oversaturation, which does not cause immediate embolism formation. Pneumatic experiments on dissolved and undissolved gas in xylem combined with gas diffusion models provide exciting opportunities to better understand the mechanisms underlying xylem embolism formation, and how plants may avoid constant hydraulic failure. We speculate that xylem sap lipids, which are essential for the production and stability of nanobubbles, together with mesoporous pit membranes and nanoporous cell walls of conduits, play a crucial role in the oversolubility of xylem sap. Our findings revealed high concentrations of gas dissolved in xylem sap, which reiterates the enigma of how plants can achieve water transport under negative pressure. In fact, the wider relevance of our work goes back to Henry Dixon (one of the originators of the cohesion-tension theory behind water transport in plants), who suggested that xylem sap may contain some unknown compounds that reduce the likelihood of embolism under negative pressure as compared to pure water (Dixon, 1914).

## Acknowledgements

We thank Andrea Huppenberger, Clara García Sanchez, and the Electron Microscopy Unit of Ulm University for preparation of the TEM samples. Mel Tyree is acknowledged for discussions about gas kinetics in xylem. S.J, L.P., L.K., and X.G. acknowledge the Deutsche Forschungsgemeinschaft (DFG, German Research Foundation; project numbers 383393940, 410768178, and 457287575). The authors acknowledge the São Paulo Research Foundation (FAPESP, Brazil) for a research grant (E.C.M, R.V.R., L.P. and M.T.M., Grant 2019/15276-8), a fellowship (L.P. and R.V.R., Grant 2017/14075-3), and a scholarship (M.T.M. and R.V.R., Grant 2018/09834-5). E.C.M., R.V.R. and JLSM are fellows of the National Council for Scientific and Technological Development (CNPq, Brazil). H.J.S. and S.J. acknowledge financial support from the National Science Foundation (IOS-1754850).

## Authors contribution

LP conceived the idea and identified the daily patterns of gas extraction. LP, RVR, SJ, and HJS developed the hypotheses and planned the experiments, which were conducted by LP, MTM, VSP, GSP, JLSM, LK, XG, and ECM, who also contributed to the hypotheses’ discussion. All authors contributed to the manuscript writing, with substantial inputs from RVR, SJ, HJS, and LP.

