## Supplementary Material for "Dynamic changes in gas solubility of xylem sap reiterate the enigma of plant water transport under negative pressure"

**Methods S1** Sap gas extraction model (SAX)

To develop the SAX model, we needed various anatomical parameters for branches of *Citrus sinensis*, a gas diffusion coefficient, and transport rate. Based on variables shown in Table 1, we calculated the mean vessel length $V_L$ (m), the cut-open vessel length ($V_{openL}$) and volume ($V_{openV}$), the total surface area occupied by intervessel pit membranes ($A_p$) following Wheeler *et al.* (2005), the number of vessels ($V_n$), the total cut-open vessels volume ($V_{open\_total}$), and the total volume of cut-open vessels plus discharge tube volume ($V_t = V_r + V_{open\_total}$), as follows:

$$V_{open_L} = \frac{V_L}{2} \tag{4}$$

$$V_{open_V} = \pi V_{open_L} \left(\frac{V_d}{2}\right)^2 \tag{5}$$

$$A_p = \pi V_d V_L F_{PM} V_F \tag{6}$$

$$V_n = V_{Lumen} \left(\frac{S_d}{V_d}\right)^2 \tag{7}$$

$$V_{open_{total}} = V_n V_{open_V} \tag{8}$$

$$V_t = V_r + V_{open_{total}} \tag{9}$$

Henry's constant ($K_H$, in bars) was estimated considering the measured water potential (Ψ) and temperature (T) during our experiments, according to Mercury et al. (2003) and Schenk et al. (2016):

$$K_H = e^{\left(\frac{a+b*\psi+c*T}{\ln(T)}\right)} \tag{10}$$

with values of *a*, *b* and *c* referring to gases shown in Table S1.

The solubility of a particular gas ($C_{s[g]}$) in xylem sap was then calculated as:

$$C_{s[g]} = \frac{P_s}{K_H} \tag{11}$$

where $P_s$ was the partial gas solubility. We considered that the atmosphere was composed of $N_2$ and Ar (78.09% and 0.93%, respectively), and variable proportions of $CO_2$ and $O_2$, varying from zero to 20.98%. For this, we used measurements of $CO_2$ concentration in *Citrus* plants, and $C_s$ was calculated as the sum of the partial solubilities.

The gas concentration in air ($C_{gas}$) at atmospheric pressure (P = 101300 Pa) was calculated as:

$$C_{gas} = \frac{P}{(RT)} \tag{12}$$

where R is the gas constant (8.3144621 J $mol^{-1}$ $K^{-1}$) and T is the air temperature (in K).

The radial diffusion at the cut-open vessel (Fig. 1b,d) was estimated as described by equations 5 to 8 in Yang *et al.* (2021). For the gas extracted axially from the sap in intact vessels, we considered the contact area between the sap and the pit membrane, and assumed that the gas diffusion coefficient in water ($D_w$) was $2*10^{-5}$ $m^2$ $s^{-1}$. Then, the axial transport rate ($k_w$) through sap at every $2.5*10^{-5}$ m ($d_w$) step from the pit membrane, was estimated over 150 seconds:

$$k_w = \frac{\Delta t A_p D_g}{d_w} \tag{13}$$

Where $\Delta t$ is the time interval of a discharge measurement, $A_p$ is the total pit area, and $D_g$ is the diffusion coefficient of gas in water. We calculated the number of mols ($\Delta n_w$) for 500 steps (index i). When we considered more than 500 steps, the amount of gas extracted was relatively constant (Fig. S2). Hence, most of the gas could be extracted from the first 500 steps, which resulted in a total distance ($d_w$ * steps) of 1.25 cm from the pit membrane:

$$\Delta n_w = k_w(C_{s,i} - C_{s,i-1}) \quad (14)$$

The gas concentration in sap ($C_s$) in each time step j was then calculated as:

$$C_{s,j} = C_{s,j-1} + (\frac{\Delta_{n,i} + \Delta_{n,i-1}}{V_v}) \quad (15)$$

where $V_v$ was the volume of a single vessel.

The gas transport constant through pit membranes ($k_a$) was estimated as:

$$k_a = \frac{\Delta t A_p D_g H^{CC}}{d_m} \quad (16)$$

where $d_m$ is the pit membrane thickness and $H^{CC}$ is the dimensionless Henry's constant, which was calculated by dividing $C_s$ by $C_{gas}$.

We simulated the gas extraction by the vacuum pressure applied to the discharge tubing (*ca.* 35 kPa of absolute pressure) as described in Yang *et al.* (2021). Then, the change in the amount of gas extracted axially ($\Delta n_a$, mol) for each measuring second at the cut-open vessel and Pneumatron discharge tube was:

$$\Delta n_a = k_a(C_{s,j} - C_{g,j} * H^{CC}) \quad (17)$$

where at time step j, $C_{s,j}$ was the total gas concentration in xylem sap, and $C_{g,j}$ was the gas concentration in the cut-open vessels and discharge tube.

Then, the total gas concentration in the cut open vessels and discharge tube ($C_g$, in mol $m^{-3}$) was calculated as the sum of the radial ($V_n\Delta_{nr}$) and axial ($V_n\Delta_n$) gas discharged divided by the total volume of the cut-open vessels and the discharge tube of the Pneumatron ($V_t = V_{open\_total} + V_r$):

$$C_g = \frac{V_n \Delta_n + V_n \Delta_{nr}}{V_t} \quad (18)$$

where $\Delta_{nr}$ was the number of mols discharged radially (see equation 8 in Yang *et al.* (2021)).

The amount of gas extracted (G, in mol $m^{-3}$) and the absolute amount of gas discharged (GD, in mol) were estimated as:

$$G = (C_{g[150s]} - C_{g[1s]}) \quad (19)$$

$$GD = G\,V_t \quad (20)$$

where $C_g$ was estimated at the first second and after 150 s of gas extraction.

*Modelled versus measured gas extraction*

We calculated the expected gas discharge ($GD_{expected}$) from the expected gas concentration in sap ($C_{s\_expected}$), using the SAX model described above and considering measurements of temperature, water potential, and $CO_2$ concentration in *Citrus* plants. We also used the slope and intercept of the linear relationship between $GD_{expected}$ and $C_{s\_expected}$ to estimate the gas concentration dissolved in xylem sap ($C_s$) from the gas discharge values measured (GD) in the experiments 3 and 4. Our results were also shown as the proportion of $C_s$ related to $C_{s\_expected}$. In addition, we checked the contribution of the radial gas extraction to the total gas extraction, and consequently the importance of the axial extraction. For this, the equations 13 to 17 were not considered.

Finally, we evaluated the contribution of water vapour to the gas discharge measurements, estimating the absolute humidity at the cut-open vessels and assuming an equilibrium between the pressure of the liquid within intact vessels and surrounding tissues, and gas inside the cut open-vessels and Pneumatron tubing. For this, we estimated the relative humidity ($U_r$) as:

$$U_r = e^{\left(\frac{\psi * V_t}{RT}\right)} \tag{21}$$

The saturated vapour pressure ($e_s$, in Pa) and the absolute humidity ($U_a$, mol $m^{-3}$) were estimated as described in equations 22 and 23 (Jarraud, 2008):

$$e_s = 614.1e^{\left(\frac{17.62T}{T+243.12}\right)} \tag{22}$$

$$U_a = U_r \frac{e_s}{RT} \tag{23}$$

Where T is the temperature in Celsius. All analyses and models were developed using R programming environment, version 3.6.3 (R Core Team, 2020). Codes are available upon request.

**Table S1** Parameters used for calculating Henry's constant ($K_H$) for four major gases (eq. 9).

| gas | *a* | *b* | *c* |
|---|---|---|---|
| Ar | 10.15092 | 0.001215 | 0.054913 |
| $N_2$ | 10.96579 | 0.001275 | 0.046013 |
| $O_2$ | 10.23061 | 0.001167 | 0.056181 |
| $CO_2$ | 6.647762 | 0.001236 | 0.099455 |

**Box. S1** Workflow of the gas extraction measurements and SAX-model, describing how the expected gas concentration in sap was compared to the estimated values from the gas discharge measured in plants of Citrus sinensis. The abbreviations are described in Methods S1, and summarized in Table 2.

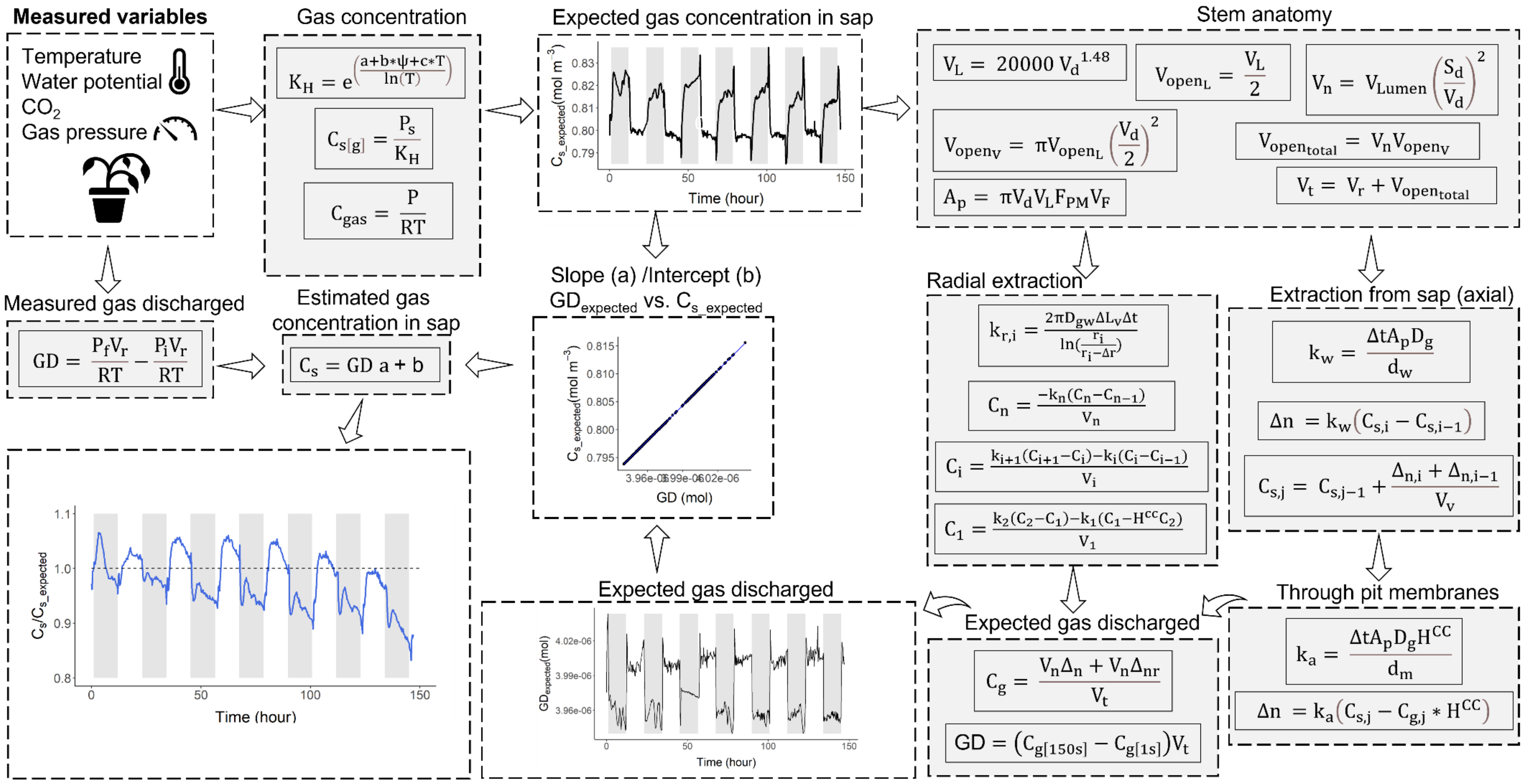

**Fig. S1** (a) Vulnerability curve of *Citrus sinensis* (data from Pereira *et al.* 2020) measured in similar plant material as used in our experiments (n = 4). The gas discharged and the xylem water potential were measured with a Pneumatron (Pereira *et al.* 2020) and a stem psychrometer (PSY1, ICT International, Armidale NSW, Australia), respectively. The continuous blue line is the sigmoidal fit and the dashed line is pointing to the water potential in which 50% of embolism was estimated from the gas discharge measurements; (b) Gas discharged related to water potential of the same data shown in (a) (black circles) compared to the *in vivo* measurements in our five experiments (green symbols). We considered 15 seconds of gas extraction.

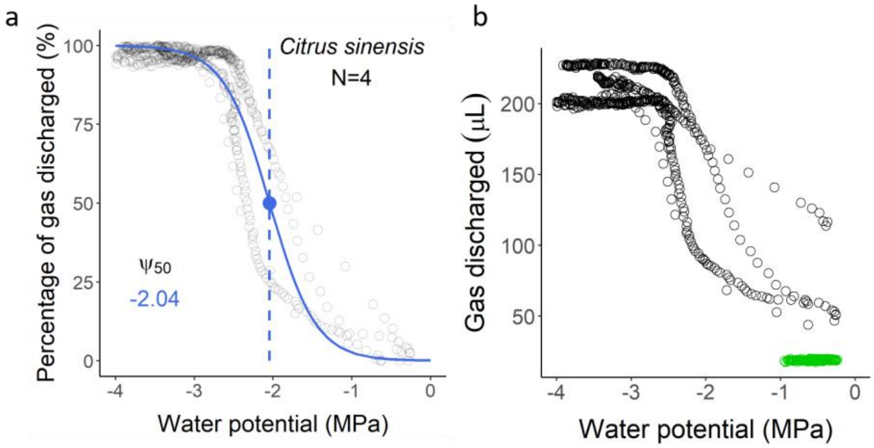

**Fig. S2** Modelled gas concentration in the Pneumatron discharge tube at the 150 s of gas extraction in function of the number of steps considered to simulate the axial gas extraction from sap in intact vessels. We considered $\Delta t = 2.5 * 10^{-5}$ s and $d_w = 2.5 * 10^{-5}$ m, which resulted in gas extraction from sap in intact vessels over a total distance ($d_w$ * steps) of 1.25 cm.

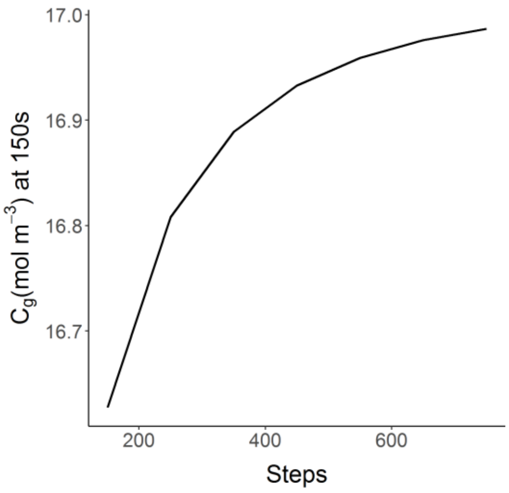

**Fig. S3** Temporal dynamics of the percentage of gas discharged related to the maximum values measure ($PGD_{max}$, green line) from xylem sap of *Citrus sinensis* trees, with reference to temperature (orange line) under 12 hours of light (white bars) and 12 hours of darkness (grey bars) over various days. Each graph represents one plant. $PGD_{max}$ decreased after five days and the daily variation stopped after ca. 13 days, reaching a minimum gas extracted of 11.2 ± 1%.

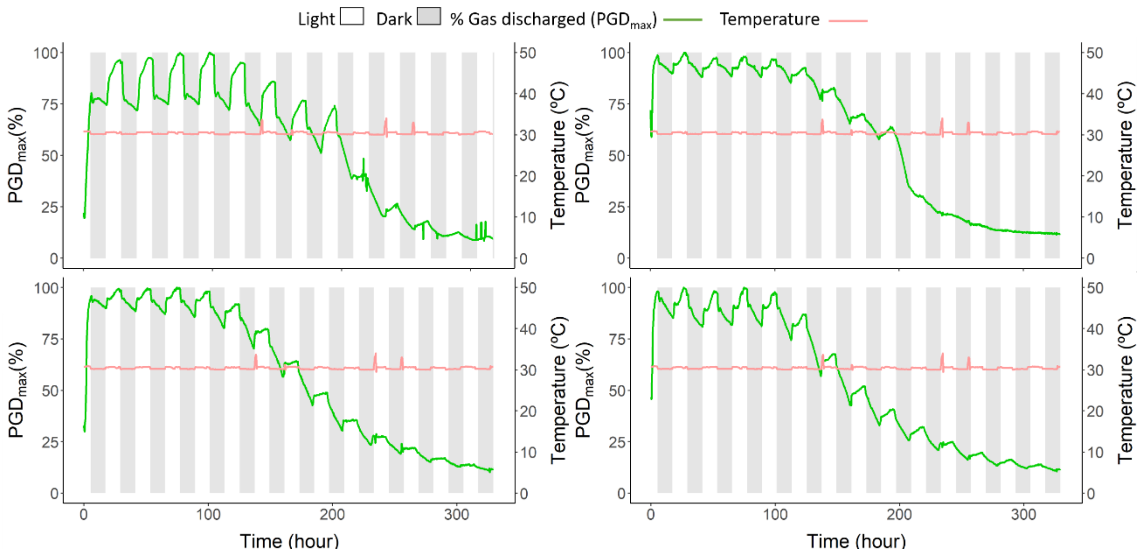

**Fig. S4** Temporal dynamics of the percentage of gas discharged (PGD, green line) from xylem sap of *Citrus sinensis* plants under varying temperature (orange line) and photoperiodicity (light = white bars, dark = grey bars) over 7 days. Box plots compare all data points from the graphs in the first column. Each graph represents a single plant. In (a) and (b), plants were submitted to 30°C or 15°C with constant light for three days, and then submitted to light or dark conditions under constant temperature (30°C) for an additional four days. The opposite sequence was applied in (c), (d), and (e).

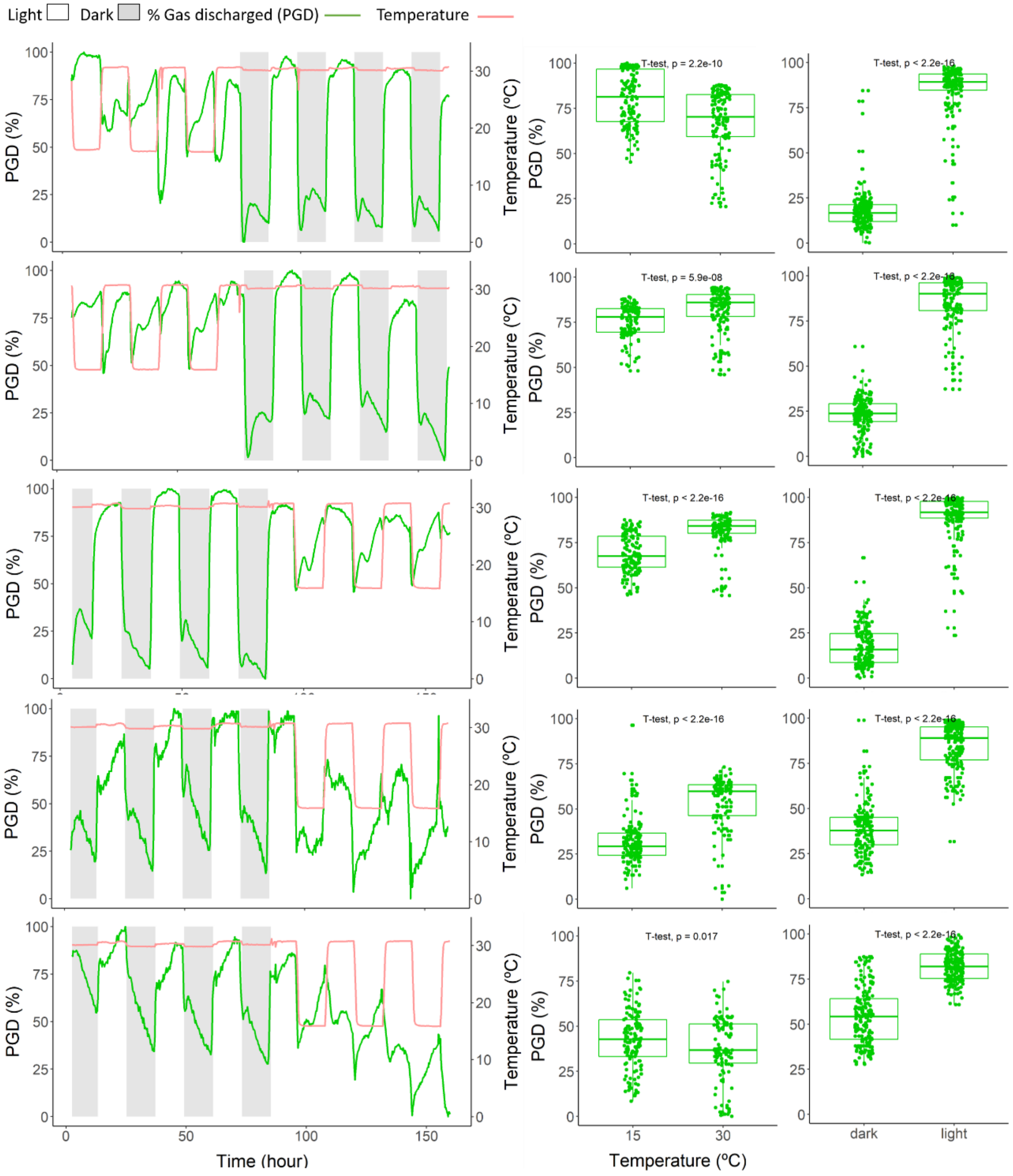

**Fig. S5** Percentage of gas discharged (PGD, green line) from xylem sap of *Citrus* sinensis plants and xylem water potential (blue line) under light (white bars = 12 hours) or dark (grey bars= 12 hours) over time. The scatter graphs correlate PDG and water potential from the graphs on the left, with the respective linear regression (in blue). Each graph represents in vivo measurements on a single branch of an individual plant.

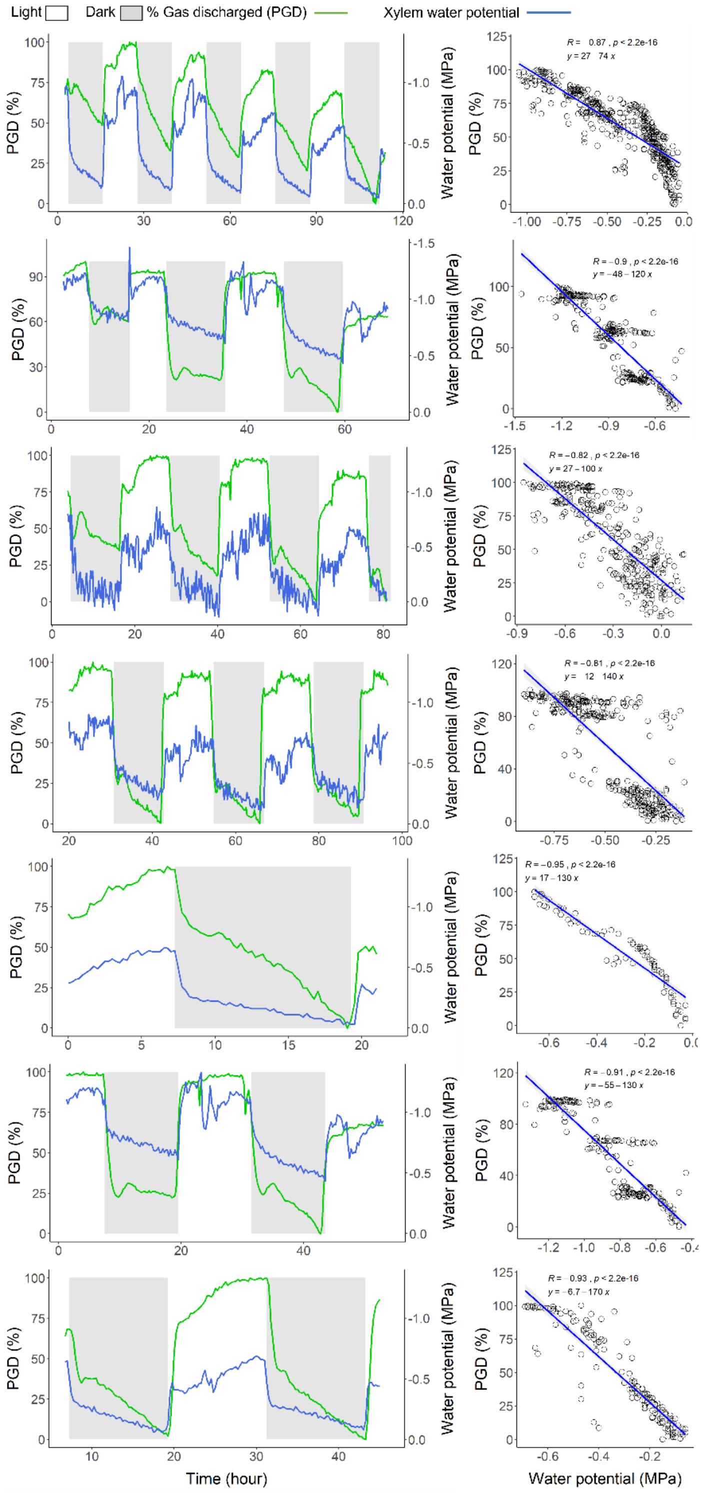

**Fig. S6** Temporal dynamics of the percentage of gas discharged (PGD, green line) and xylem water potential (blue line) as shown in the left graphs. The graphs on the right correspond to the same respective plants, showing the measured $CO_2$ concentration before measurements started (black line), after one second of gas extraction (green line), and after 150 seconds of extraction (blue line) . Shown are also the air temperature (orange line), periods of 16 hours of light (white bars) or 8 hours of darkness (grey bars). The measurements run over 6 to 7 days. All the leaves were removed, and the stems covered with aluminum foil after 2 to 3 days (vertical dashed lines) to stop transpiration and gas exchange as much as possible.

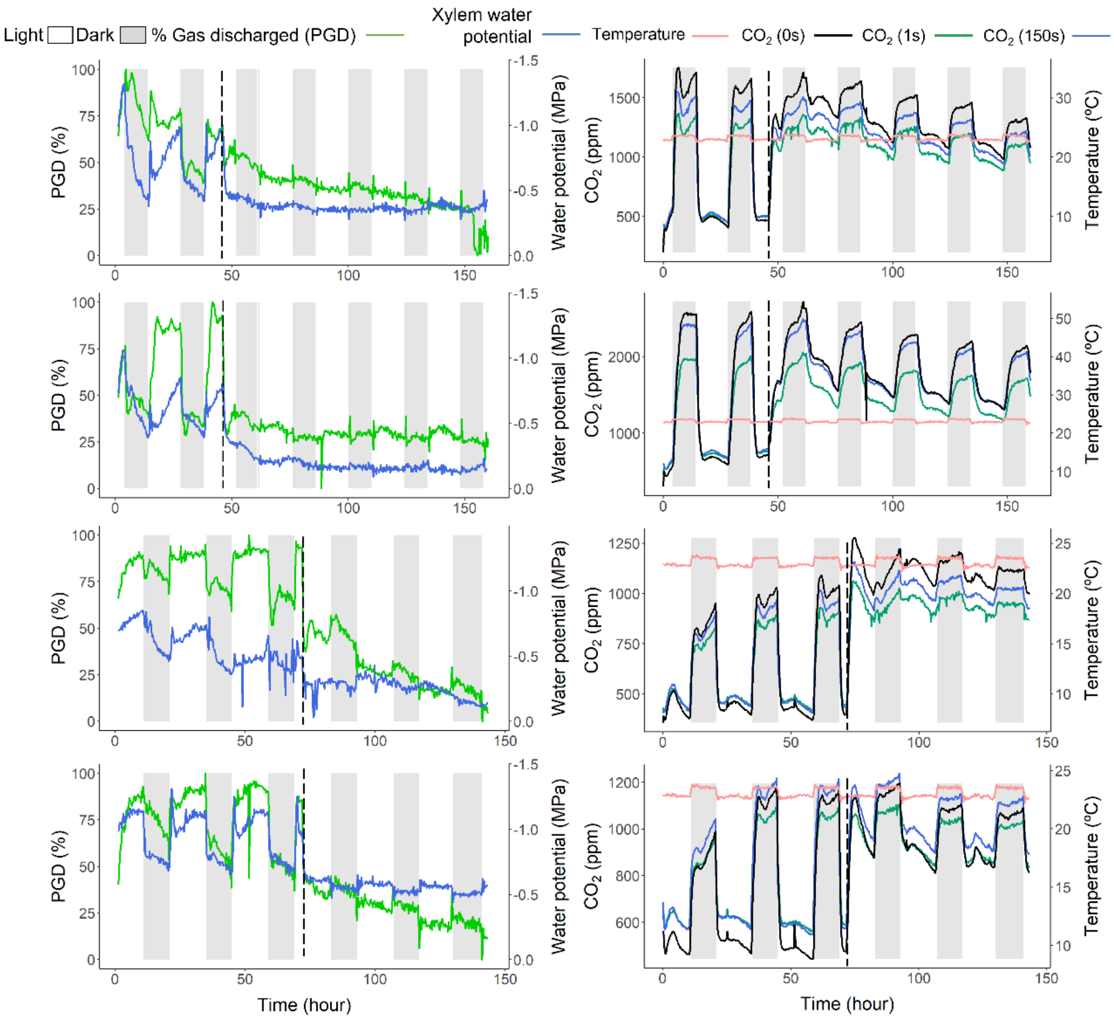

**Fig. S7** Percentage of gas discharged (PGD, green lines) from xylem sap of *Citrus sinensis* plants and transpiration (blue lines) under 12 hours of light (white bars) or 12 hours of darkness (grey bars) over time. The scatter graph on the right correlates PGD and water potential from the graphs on the left, with the linear regressions shown (in blue).

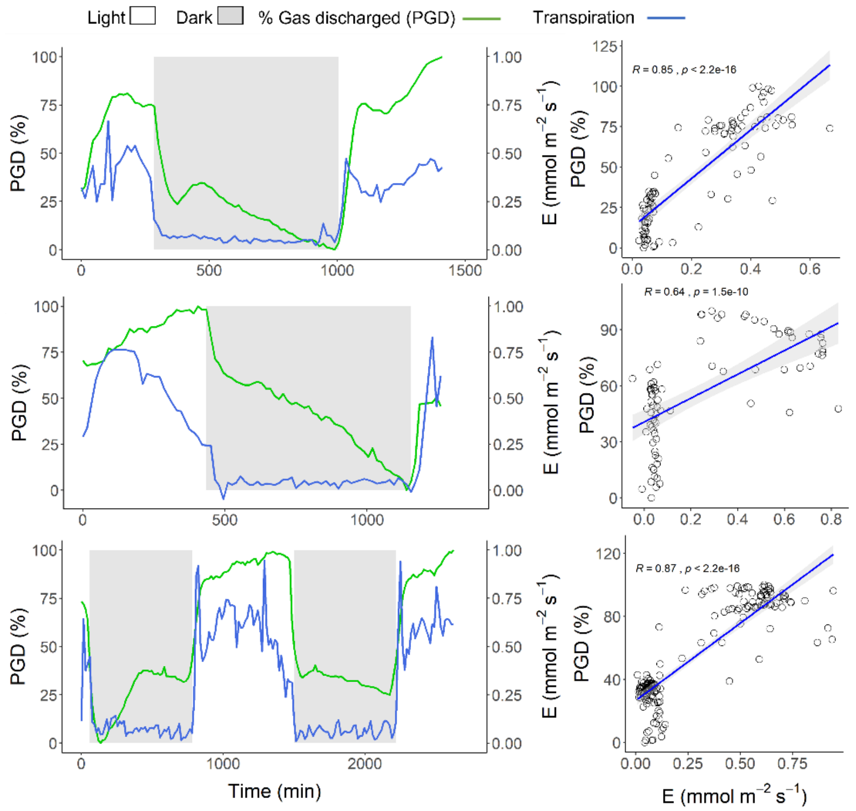

**Fig. S8** Percentage of gas discharged (PGD, green lines) from xylem sap of *Citrus sinensis* plants, and the photosynthetic rate (blue lines) under 12 hours of light (white bars) or 12 hours of darkness (grey bars) over time. The scatter graph on the right correlates PGD and water potential from the graphs on the left, with the respective linear regressions (in blue).

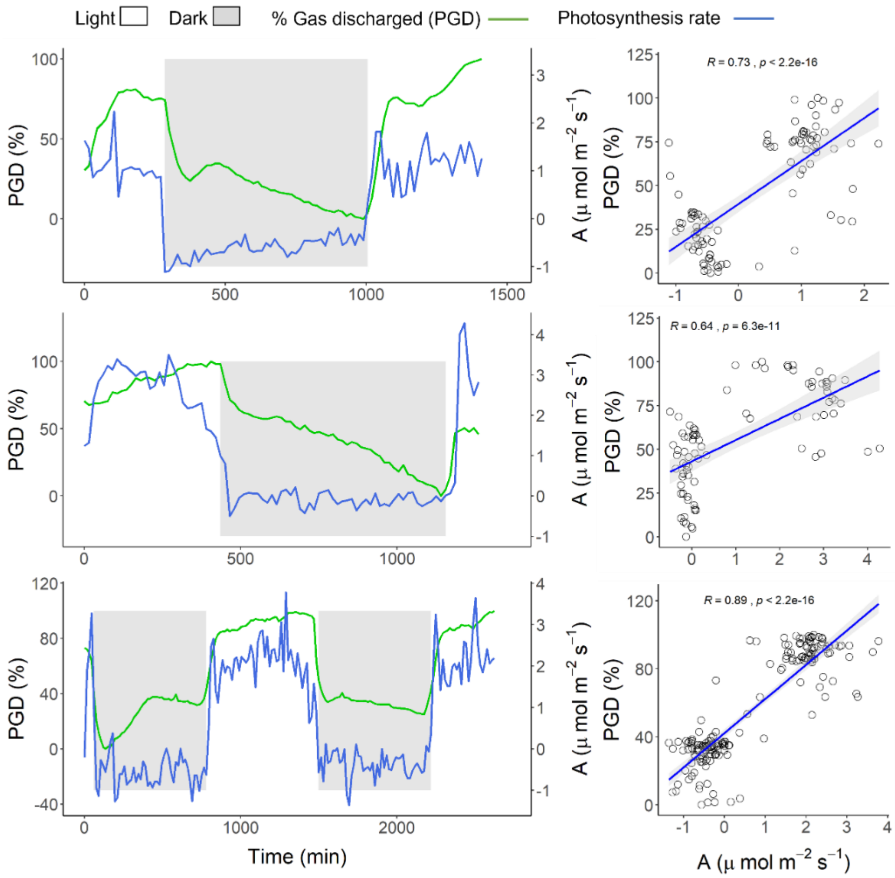

**Fig. S9** Percentage of gas discharged (PGD, green lines) from xylem sap of *Citrus sinensis* plants, and stomatal conductance (blue lines) under light (white bars = 12 hours) or dark (grey bars = 12 hours) over time. The scatter graph on the right correlates PGD and water potential from the graphs on the left, with the respective linear regressions (in blue).

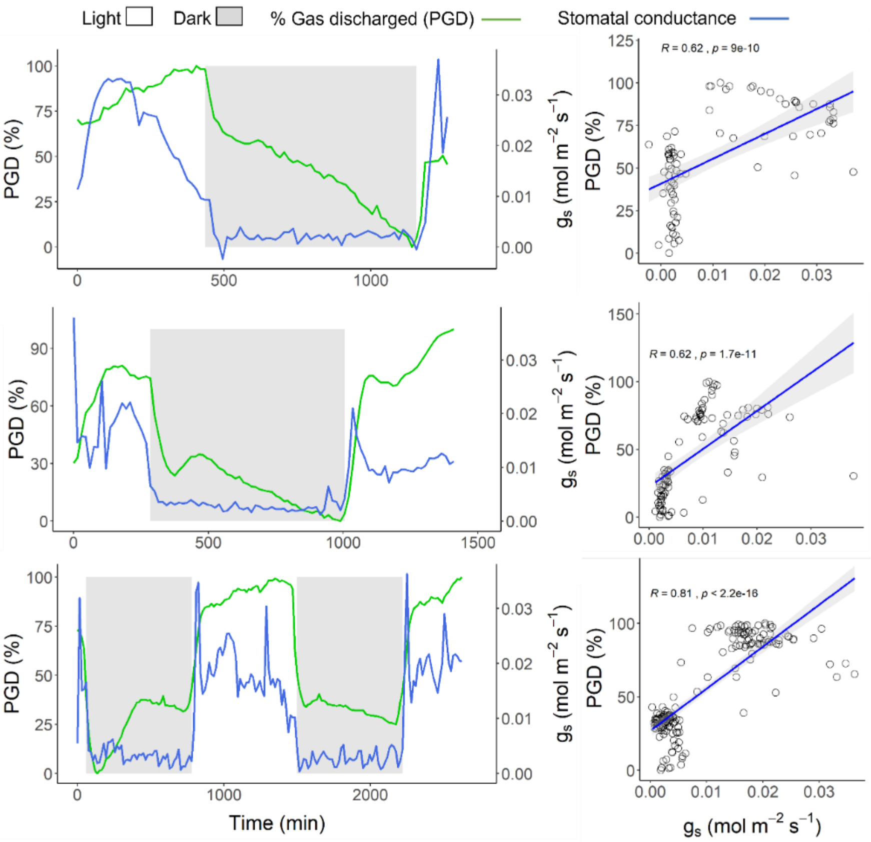

**Fig. S10** Percentage of gas discharged (PGD, green lines) from xylem sap of *Citrus sinensis* plants and internal carbon concentrations in leaves (blue lines) under light (white bars = 12 hours) or dark (grey bars = 12 hours) over time. The scatter graph on the right correlates PGD and water potential from the graphs on the left, with the respective linear regressions (in blue).

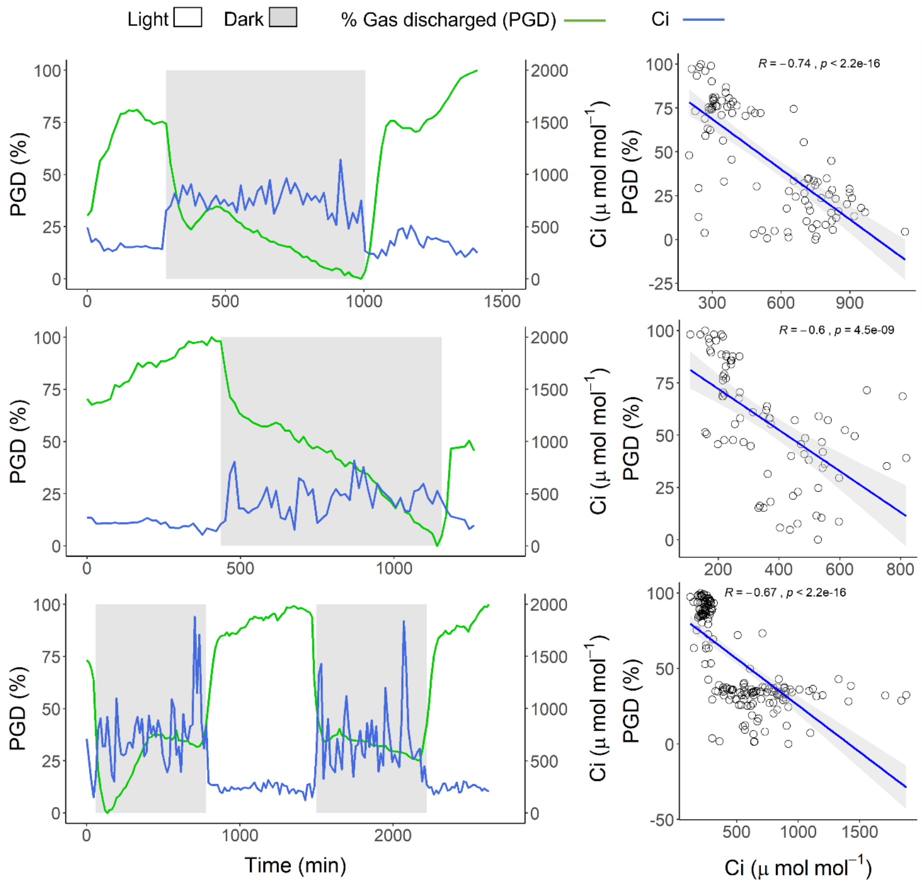

**Fig. S11** (a) Temporal dynamics of the percentage of gas discharged (PGD, green points) under light (white bars) or dark (grey bars) conditions over 5-8 days. The vertical dashed line indicates when a short segment of the branch was cut and sealed with glue on the distal side, keeping less than 2 cm connected to the Pneumatron. Box plots in (b) compare all data points shown in (a), before (total amount of gas discharged), and after reducing the stem to less than 2 cm, which allowed estimating the background gas from xylem tissue (labeled as "leakage").

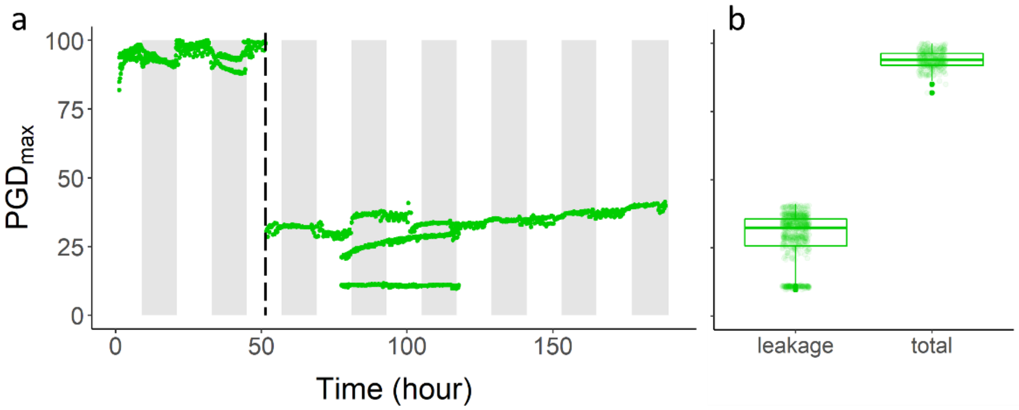

**Fig. S12** Probability distribution of vessel length (a), vessel diameter (b) and vessel lumen fraction (c) measured in six branch samples of *Citrus sinensis*. In (a), the calculated mode and mean (black and blue values and dashed lines, respectively) and the standard deviation (the range in red) are shown.

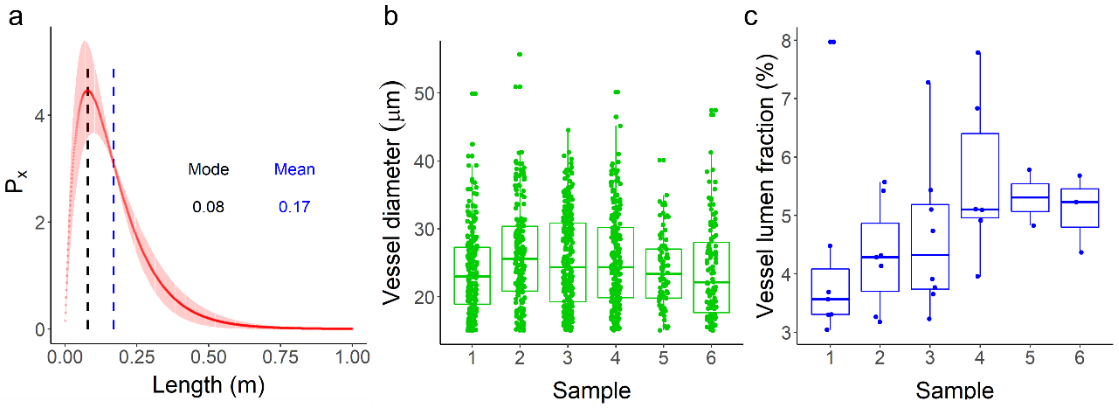

**Fig. S13** Temporal dynamics of the modelled gas concentration in xylem sap ($C_s$) in relation to the expected gas concentration ($C_{s\_expected}$) in a well-irrigated *Citrus sinensis* tree exposed to 12 hours of light (white bars) and 12 hours of darkness (grey bars), considering (a) the mean and the standard deviation of the anatomical parameters for which the model was more sensitive to predict gas concentration. The signals - and + indicate mean values ±SD for the following parameters: vessel diameter ($V_d$), intervessel fraction ($V_F$), and vessel lumen fraction ($V_{lumen}$), and different colours indicate the results considering all the possible combinations among these parameters. The dotted line represents the $C_s/C_{s\_expected}$ ratio when the mean values were considered. In (b) the ±SD for $V_d$ (red), $V_F$ (green), or $V_{lumen}$ (blue), when using the mean values for the other two parameters. The horizontal dashed lines in (a) and (b) represents the 1:1 line.

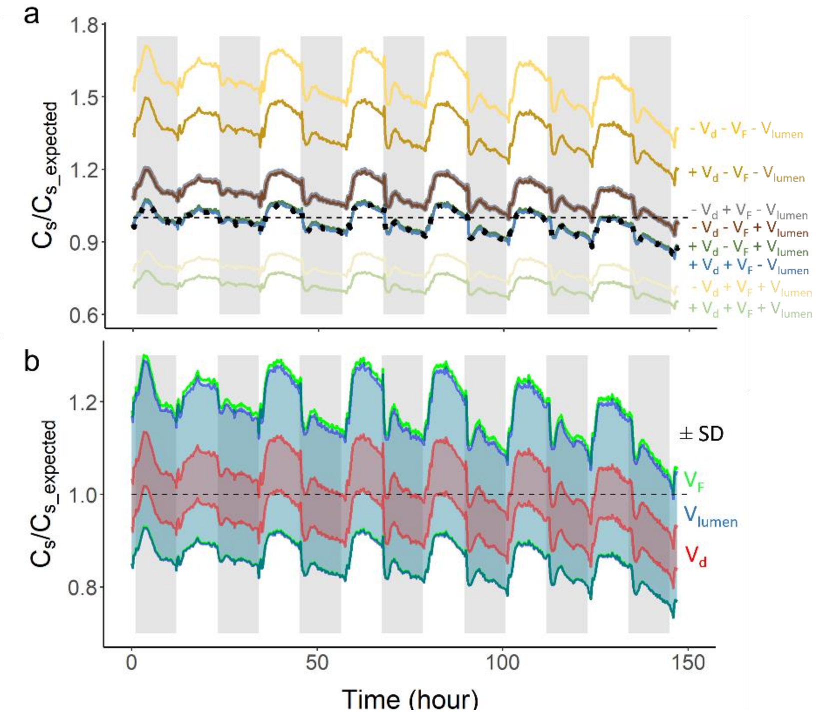
